# Neuro-transcriptomic signatures for mood disorder morbidity and suicide mortality

**DOI:** 10.1101/762492

**Authors:** Mbemba Jabbi, Dhivya Arasappan, Simon B. Eickhoff, Stephen M. Strakowski, Charles B. Nemeroff, Hans A. Hofmann

## Abstract

Suicidal behaviors are strongly linked with mood disorders, but the specific neurobiological and functional gene-expression correlates for this linkage remain elusive. We performed neuroimaging-guided RNA-sequencing in two studies to test the hypothesis that imaging-localized gray matter volume (GMV) loss in mood disorders, harbors gene-expression changes associated with disease morbidity and related suicide mortality in an independent postmortem cohort. To do so, first, we conducted study 1 using an anatomical likelihood estimation (ALE) MRI meta-analysis including a total of 47 voxel-based morphometry (VBM) publications (i.e. 26 control>major depressive disorder (MDD) studies, and 21 control>bipolar disorder (BD) studies) in 2387 (living) participants. Study 1 meta-analysis identified a selective anterior insula cortex (AIC) GMV loss in mood disorders. We then used this results to guide study 2 *postmortem* tissue dissection and RNA-Sequencing of 100 independent donor brain samples with a life-time history of MDD (N=30), BD (N=37) and control (N=33). In study 2, exploratory factor-analysis identified a higher-order factor representing number of Axis-1 diagnoses (e.g. substance use disorders/psychosis/anxiety, etc.), referred to here as *morbidity* and suicide-completion referred to as *mortality*. Comparisons of case-vs-control, and factor-analysis defined higher-order-factor contrast variables revealed that the imaging-identified AIC GMV loss sub-region harbors differential gene-expression changes in high morbidity-&-mortality *versus* low morbidity-&-mortality cohorts in immune, inflammasome, and neurodevelopmental pathways. Weighted gene co-expression network analysis further identified co-activated gene modules for psychiatric morbidity and mortality outcomes. These results provide evidence that AIC anatomical signature for mood disorders are possible correlates for gene-expression abnormalities in mood morbidity and suicide mortality.

## INTRODUCTION

Major depressive disorder and bipolar disorder ⎯ here together referred to as mood disorders, are the third leading cause of the global disease burden (Collins et al. 2011; Murray et al. 2012). Mood disorders account for the majority of completed suicides (Waern et al. 2002; Marangell et al. 2006) and they were linked to ∼47,000 suicides in the United States in 2017 alone (American Foundation for Suicide Prevention, 2019). However, the convergent neurobiological basis for mood symptoms/syndromes and suicide is unknown, limiting advances in developing novel interventions.

Neuroimaging studies have identified reduction in gray matter volume (GMV) in the anterior insular cortex (AIC) and anterior cingulate cortex (ACC) in association with diagnosis of psychiatric disorder in general (Goodkind et al. 2015), and the regional GMV volume reductions in these AIC and ACC network have been especially implicated in mood disorder diagnoses in particular (Wise et al. 2017). Neurobiological integrity of the right AIC is shown to (a) predict mood diagnostic severity (Hatton et al. 2012), (b) modulate subjective responses to distress, pain, and psychosocial adversity (Wager et al. 2013; Eisenberger 2015), (c) regulate affective interoception (Craig 2009; Slavich et al. 2010), (d) associate with stress-related inflammatory markers (Khalsa et al. 2018), and (e) predict psycho- and pharmaco-therapeutic efficacy in mood disorders (McGrath et al. 2013). AIC-ACC functional connectivity during affective processing differentiated mood disorder suicide-attempters from non-attempters (Pan et al. 2013). Furthermore, abnormalities in AIC volume and synaptic abnormalities are linked to suicidal-behavior in mood disorder (Wagner et al. 2012; Mathew et al. 2013). AIC response to stress is shown to impact hypothalamic-pituitary-adrenal (HPA) axis-driven inflammatory responses (Khalsa et al. 2013), which may serve to exacerbate mood disorder associated psychiatric morbidity and suicidal-behavior (Oquendo et al. 2014; Wohleb et al. 2016). Although a preponderance of evidence supports abnormal AIC integrity in psychosocial distress (Schneidman 1998; Mee et al. 2011; Wager et al. 2013) and mood/comorbid psychiatric symptoms, an underlying functional genetic contribution in terms of functional gene-expression changes for these abnormalities remains largely unknown.

The lack of a well-defined relationship between aberrant brain structure and function with underlying molecular changes within this brain region is an impediment to understanding pathophysiology. Moreover, evidence for shared genetic mechanisms underlying psychiatric diagnoses (Brainstorm consortium, Anttila et al. 2018) is not well-integrated with brain imaging correlates of psychiatric disease-morbidity and specific behaviors, in this case, suicide. In the present study, MRI meta-analysis was used to test the hypothesis that reduced AIC volume will be the most prominent neuroanatomical signature for mood disorder diagnoses. We confirmed this hypothesis with our meta-analytic findings in study 1 and then used this anatomical hallmark to guide dissection of *postmortem* brain tissue for analyses of molecular/gene-expression signatures that could pave the way for precision profiling of gene functions underlying mood symptoms across diagnoses in clinically-relevant brain sub-regions in study 2. This approach enabled us to further test the hypothesis that the voxel-based morphometry (VBM) imaging meta-analysis defined in study 1 will harbor postmortem gene-expression signatures for psychiatric disease morbidity and related suicide-mortality in postmortem mood disorder brains, thereby providing a neurobiological framework for characterizing convergent neural-and-gene expression signatures for behavioral brain diseases.

## METHODS

### PARTICIPANTS

The imaging meta-analysis provided a consolidation of the current mood disorder VBM work by quantitatively integrating published results of volumetric comparisons of interest between controls>mood disorder participants, or correlations of volumetric measures with mood disorder symptom-specific measures. In total, 47 previous VBM publications consisting of volumetric comparisons or experimental contrasts assessing GMV reductions in healthy controls > major depression from 26 independent publications (demographics in **Table 1A**); and volumetric comparisons or experimental contrasts assessing GMV reductions in healthy controls > bipolar disorder from 21 independent publications (demographics in **Table 1A**). Only publications with more than 20 subjects per comparison (i.e. publications with more than 10 subjects per each comparison group of major depressive disorder<controls, or bipolar disorder< controls) were selected based on the rationale that comparison group averages of less than 10 subjects per diagnostic or control group is likely underpowered in VBM studies. Together, the meta-analysis included 2387 imaging participants.

**Table 1.**
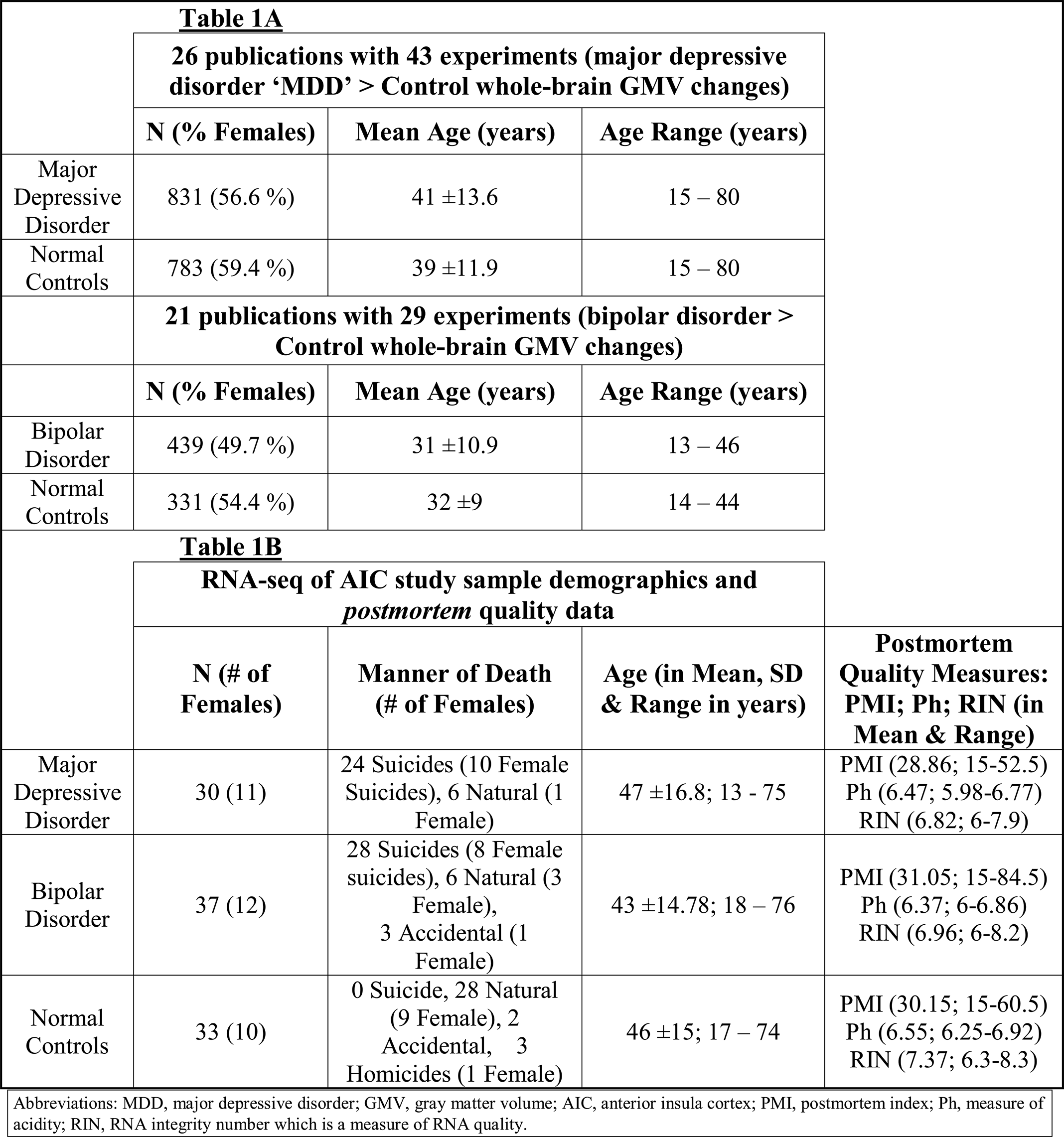

The 47 publications included in our meta-analysis, as well as relevant publications that ended up not being included based on the above-mentioned inclusion criterion, are listed in **Supplementary Table 1**. While 3 of the 72 experimental contrasts included in the meta-analysis assessed suicidal behavior in relation to volumetric changes in mood disorder, suicidal phenotypes was not a specific selection criteria for study inclusion as there were very few studies in the BrainMap database that specifically assessed the relationship between suicidal phenotypes and brain volume. In study 2, RNA samples were extracted from the anatomical likelihood estimation (ALE) defined AIC sub-regional (identified in study 1) *postmortem* tissue of 100 donors from NIMH brain bank (postmortem technical/qualitative variables in **Table 1B**).

## DESIGN

Based on the principle that neural structure subserves functional control of complex behavioral repertoires (Koechlin 2016), we localized the structural brain signature for mood disorders across samples and methods in study 1. Experiments of GMV changes associated with mood disorder diagnoses in defined stereotaxic space were included for analysis of localized GMV changes across studies in major depressive disorder, bipolar disorder, and major depressive disorder and bipolar disorder versus (vs) controls using the well-established ALE algorithm (Eickhoff et al. 2009). This signature guided localized anatomical-dissection of *postmortem* tissue for whole-transcriptome characterization of differential gene-expression and weighted gene co-expression network analysis (WGCNA).

### NEUROIMAGING VBM META-ANALYSIS

For the VBM meta-analysis, results of anatomical changes reported in coordinate space using the Montreal Neurological Institute (MNI) or Talairach coordinates of 3-dimensional brain space were included from 26 publications examining *major depressive disorder < controls* whole-brain GMV changes; 21 publications examining *bipolar disorder < controls* GMV changes; and we combined the coordinates for the 26 publications examining *major depressive disorder < controls* and the 21 publications examining *bipolar disorder < controls* GMV changes for the pooled *mood disorder < controls* comparison. In a nutshell, the ALE approach applied here utilized in the VBM meta-analysis by modeling the spatial uncertainty associated with each reported 3-dimensional brain location of significant between-group differences in GMV changes (Eickhoff et al. 2009). The meta-analysis therefore performed a rigorous ALE by comparatively assessing GMV changes in (i) *major depressive disorder < controls*, (ii) *bipolar disorder < controls*, and (iii) *mood disorders < controls*. Here, models the spatial uncertainty associated with each reported location of significant between-group differences in GMV changes (Eickhoff et al., 2009; 2012) and performed ALE assessment of GMV changes in (i) major depressive disorder<controls, (ii) bipolar disorder<controls, and (iii) mood disorders in general (i.e., pooled across major depressive disorder and bipolar disorder) vs controls.

### BRAIN DISSECTION, RNA-EXTRACTION and SEQUENCING

#### Brain Dissection

The NIMH Human Brain Collection Core (HBCC) provided the *Postmortem* samples for which informed consents are acquired according to NIH IRB guidelines. Clinical characterization, neuropathology screening, and toxicology analyses followed previous protocols (Martin et al. 2006). The region of interest targeted for dissection was defined as portion of right AIC encompassing the identified reduced GMV in the completed meta-analysis (**Fig. 1A**) by the authors. Electronic image slices of the imaging-defined GMV loss volumes are shared with the HBCC neuropathologist who used these images to guide dissection of coronal tissue slabs of each postmortem donor brain at the NIH clinical center. The dissected regional volume corresponded to the anterior portion of the insula where the caudate and putamen are approximately equal in size and tissue was dissected from this section for each brain for use in study 2 RNA-sequencing (**Fig. 1B-C**).

**Figure 1.**
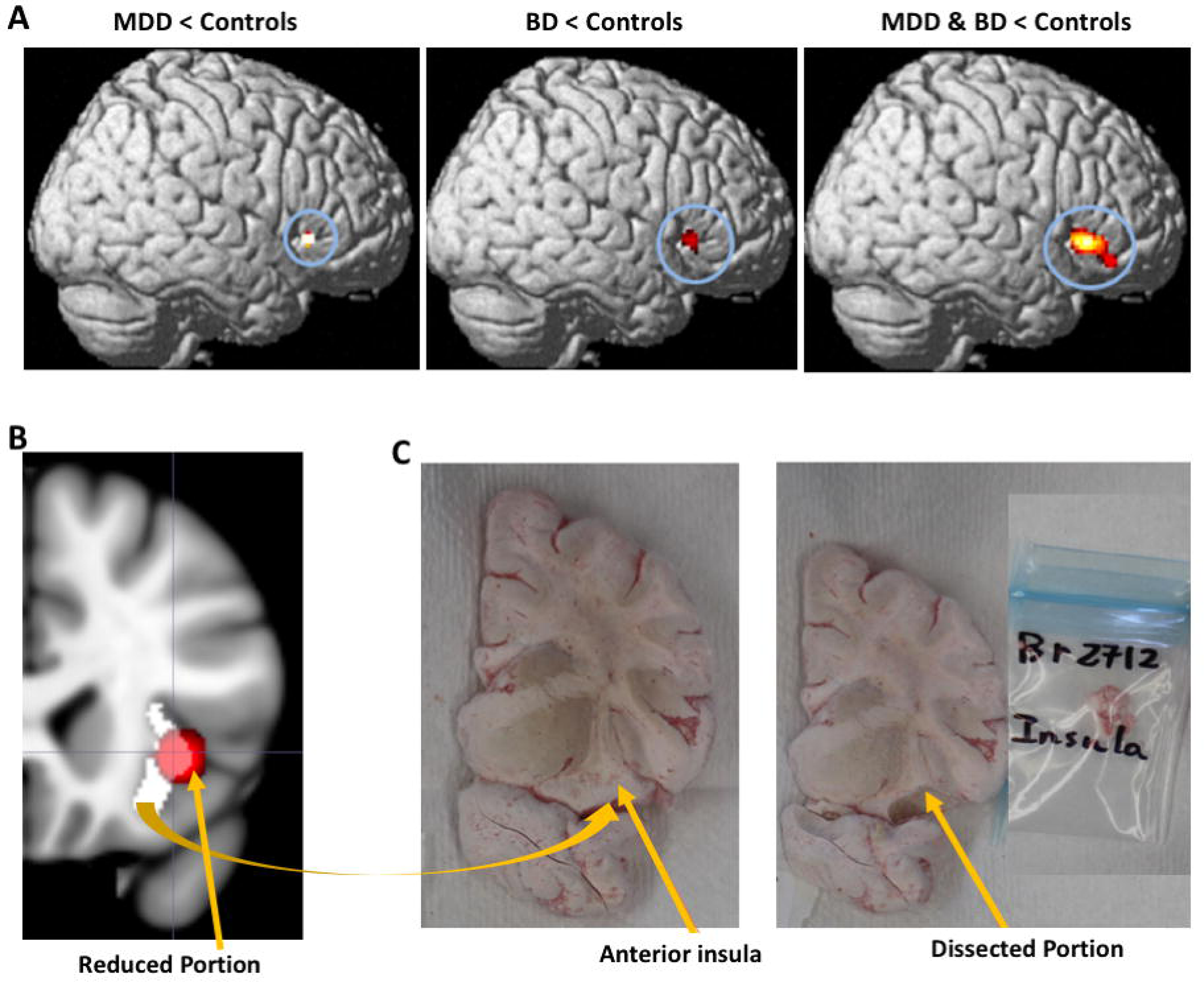
Mood disorder brain structural signature derived sub-regional dissection. A) shows reduced right AIC gray matter volume (in living) major depressive disorder<controls, bipolar disorder<controls, and the pooled mood disorder (major depressive disorder and bipolar disorder)<controls. B) shows the anatomical space demarcation of the entire right AIC (white region) and the reduced mood disorder signature (red region) and the estimated overlap (red overlapping white) between the reduced region and the AIC proper region that was targeted/dissected in the *postmortem* sample. C) *Postmortem* targeted region and the dissected portion, and dissected tissue (in labeled package).

#### RNA-Extraction

As additional service, the HBCC further pulverized all dissected tissues separately and aliquoted 50mg from each sample for standardized total RNA processing. Specifically, RNeasy Lipid Tissue Mini Kit (50) was used for RNA purification using the 50 RNeasy Mini Spin Columns, Collection Tubes (1.5 ml and 2 ml), QIAzol Lysis Reagent, RNase-free Reagents and Buffers kit from Qiagen. DNase treatment was applied to the purified RNA using Qiagen RNase-Free DNase Set (50) kit consisting of 1500 Kunitz units RNase-free DNase I, RNase-free Buffer RDD, and RNase-free water for 50 RNA minipreps. After DNAse treatment, the purified RNA from the pulverized AIC tissue sections were used to determine RNA quality as measured in RNA integrity number (RIN) values using Agilent 6000 RNA Nano Kit consisting of the microfluidic chips, Agilent 6000 RNA Nano ladder and reagents on Agilent 2100 Bioanalyzer. Samples with RIN < 6 were excluded and the 100 samples meeting inclusion were shipped directly from the NIMH HBCC core to the Genome Sequencing and Analysis Facility (GSAF: https://wikis.utexas.edu/display/GSAF/Home+Page) at the University of Texas, Austin, USA for RNA-sequencing.

#### Illumina-Sequencing, Read-Mapping and Gene-Quantification

Total RNA was extracted and only samples with RNA integrity numbers (RIN values) greater than 6 as confirmed using the Agilent Bioanayler were used for library preparation. First. Ribosomal RNA was depleted using RiboMinus Eukaryote kit from Life Technologies (Foster City, CA, USA) for RNA-Seq and confirmed using an Agilent Technologies’ Bioanalyzer (Santa Clara, CA, USA). mRNA selection was completed using the Poly(A) purist kit from Thermofisher and paired-end libraries with average insert sizes of 200bp were obtained using NEBNext Ultra II Directional RNAs Library Prep kit from New England BioLabs. All 100 samples were processed and then sequenced on the Illumina HiSeq 4000, PE150, at the Genome Sequencing and Analysis Facility (GSAF: https://wikis.utexas.edu/display/GSAF/Home+Page) at UT Austin, USA.

30 million paired-end reads per sample (150 base pairs in length) were generated by sequencing runs of 4 samples per lane of the sequencer. Sequenced reads were assessed for quality with Fastqc to specifically assess sequencing reads for median base quality, average base quality, sequence duplication, over-represented sequences and adapter contamination (Andrews 2010). The reads were pseudo-aligned to the human reference transcriptome (GRCh38-gencode) using kallisto (Kallisto, 2019), and gene-level abundances were obtained. The abundances were normalized using DESeq2, and transformed with variance stabilizing transformation (defined here as a transformation or normalization technique that seeks to create more homoscedasticity, and thereby having a closer to constant variance in the dataset regardless of the mean expression value). Principal Component Analysis was performed using 25% of the highest variance genes in order to look at the underlying structure of the data and to identify the largest sources of variance and thereby detect possible outliers.

### STATISTICAL ANALYSIS

#### VBM Meta-analysis

Convergence across the findings reported in previous VBM studies was assessed using ALE, which in brief consists of first modelling the spatial uncertainty associated with each reported location for significant between-group differences (Eickhoff et al. 2009; Turkeltaub et al. 2012).

The ALE method therefore computes the convergence across experiments by the union of the ensuing probabilistic model relative to a null-distribution reflecting a random spatial association between the findings of different experiments (Eickhoff et al. 2012). Finally, statistical inference for a whole-brain corrected significance level of p<0.001 used the threshold-free cluster enhancement method (Smith & Nichols 2009). We performed ALE separately focusing on GMV changes in controls>major depressive disorder, controls>bipolar disorder, and controls>mood disorders in general (i.e., pooled across major depressive disorder and bipolar disorder).

#### *Postmortem* variable factor-analysis

The *postmortem* variables included *mood disorder-diagnoses; # of lifetime-Axis-I diagnostic-occurrences* (e.g. Axis-I-loading of the number of comorbid disorders such as (poly)-substance use disorders, psychosis, anxiety, eating disorders, etc.); # of *lifetime-Axis-III diagnoses* (e.g. medical conditions such as diabetes, cancer, cardiovascular disease, etc.)*; manner of death* (e.g. natural, suicides, homicides or accidents) and *cause of death* as specified by the medical examiner reports (e.g. blunt force trauma to the chest, gunshot, motor vehicle accident, drowning, hanging, etc.)*; demographics* (race, age at death, sex, years of education, number of children/fecundity, and marital records); *technical variables* (brain-weight, postmortem-index, pH, and RIN-values); and *toxicology* (blood alcohol/blood narcotics levels).

We applied Principal Axis Factoring using the Oblimin Rotation with Kaizer Normalization (Costello & Osborne 2005) to identify higher-order factors explaining the differences in *postmortem* variables and included those variables with communalities of ≥ 0.5.

#### Differential Gene Expression Analysis

Just like in the imaging meta-analytic ALE, we compared AIC gene expression profiles by first conducting simple comparisons across controls>major depressive disorder, and controls>bipolar disorder.

Our factor analysis revealed that cumulatively, *mood disorder-diagnoses, Axis I diagnostic-load,* and *manner of death* together explains variability in <u>psychiatric disease morbidity</u> (data on any Axis-III/medical morbidity like cardiovascular disorders or cancer have not been accounted for in our analysis), and <u>suicide mortality</u> (i.e. mortality or completion of suicide) as a higher-order factor. The high vs low scores of this factor were determined by binning the factor scores into ≤ 0.82 for <u>low</u> and ≥ 0.82 for <u>high</u> *psychiatric morbidity-&-mortality* scores using a split-half method of dividing the maximum score across groups by 2 (i.e. 1.64/2). Specifically, we applied a T-test by binning samples into two comparison groups for profiling gene-expression between *1*) <u>high vs low</u> psychiatric morbidity-&-suicide-mortality across major depressed and control groups (i.e. major depressed with high scores vs major depressed and all controls with low scores); *2*) high vs low *psychiatric morbidity-&-suicide-mortality* scores across bipolar and control groups (i.e. bipolar with high scores vs bipolar and all controls with low scores); *3*) high vs low *psychiatric morbidity-&-suicide-mortality* scores across all groups (i.e. bipolar and major depressed with high scores vs bipolar, major depressed and all controls with low scores). Differential gene-expression between samples differing in *psychiatric morbidity-&-suicide-mortality* status was assessed based on the negative binomial distribution for modeled gene counts using DESeq2 (Anders 7 Huber 2010). RIN-values were included in the DESeq2 design matrix as a covariate to control for its potentially confounding effects.

Our factor-analysis yielded higher-order factor representing *psychiatric morbidity-&-mortality* including measures of suicide completion. In addition, our mood disorder samples have a substantially high incidence of completed suicides (i.e. 24 suicides vs 6 non-suicides out of 30 MDD and 28 suicides vs 9 non-suicides out of 37 bipolar samples). As such, we specifically examined the relationship between suicide completion and differential gene expression profiles by pooling non-suicide cases with controls. To do this, we applied a T-test to first compare **a**) gene expression difference in major depressive disorder death by suicides (n=24) vs. major depressive disorder non-suicidal deaths (n=6) & controls (n=33); and **b**) gene expression differences in bipolar disorder death by suicides (n=28) vs. bipolar disorder non-suicidal deaths (n=9) & controls (n=33); and finally compared **c**) the pooled mood disorder suicide vs. non-suicide (i.e. major depressive disorder and bipolar disorder death by suicides (n=52) vs. major depressive disorder and bipolar disorder non-suicidal deaths (n=15) without including the controls. Comparison **c** is in line with Pantazatos et al. 2017, whose results reported comparisons of major depressive disorder suicide ‘i.e. MDD-S’ vs. major depressive disorder non-suicides and controls ‘i.e. MDD+CON’ to estimate the effects of suicide. Controls were omitted in our last comparison (i.e. comparison **c**) to examine the existence of gene expression profiles that might be specifically linked to suicide completion vs non-suicide deaths in persons diagnosed with mood and comorbid psychiatric disorders. Only genes with corrected p-value (after benjamini-hochberg multiple testing correction) ≤ 0.1 and absolute fold changes 1.5 are ≥ reported as significantly differentially expressed. Pathways and gene-ontology (GO) terms enriched in these genes were identified using Enrichr (Chen et al. 2013; Kuleshov et al. 2016).

#### Weighted Gene Co-Expression Network Analysis (WGCNA)

Scale-free co-expression networks were constructed with gene-abundances using the WGCNA package in R (Langfelder & Horvath 2008). WGCNA provides a global perspective and allows identification of co-expressed gene-modules. It avoids relying on arbitrary-cutoffs involved in selecting differentially-expressed genes and instead identifies a group of genes that are changing in the same direction and magnitude, even if these changes are smaller in magnitude. WGCNA thereby identifies genes that are potentially co-regulated or belong to the same functional pathway, using a dynamic tree-cutting algorithm based on hierarchical clustering (i.e. minimum module size=30). To identify co-expressed gene modules of interest, we incorporated covariate information and selected those co-expressed gene modules correlating significantly with *diagnostic, and suicide-linked variables*. Driver genes (i.e. genes within co-expressed gene modules whose singular expression patterns are similar to the overall expression profile of the entire co-expressed modules) within these modules were used to identify pathobiological functions associated with each module.

## RESULTS

### Identification of a Mood Disorder Neuroanatomical Signature in Living Brains

The study 1 VBM meta-analysis (N=2387) revealed reduced GMV in the right AIC in mood disorders (p<0.0001 corrected) (**Fig 1A**) consistent across both major depressive disorder and bipolar disorder, since major depressive disorder and bipolar disorder groups did not differ significantly (**Fig 1A**). The localized reduced AIC neuroanatomical-signature for mood disorders was manually segmented in ITKSNAP (http://www.itksnap.org/pmwiki/pmwiki.php) (**Fig 1B**) and the segmented volume guided *postmortem* dissection of tissue used in RNA-seq characterization of gene-expression (**Fig 1C**).

### *Postmortem* Group Differences and Factor Analysis

Using principal axis factoring of the postmortem variables in study 2, we found three factors together cumulative explaining 42.22% of the variance (i.e. 17.052% of the variance was explained by <u>demographics and medical diagnostic status [</u>with high communalities loading with *number of children* (.643), *marital status* (.834), lifetime *axis-III medical diagnostic status* (.635), and *age at death* (.584)]; whereas 16.27% of the variance was explained by psychiatric disease morbidity and mortality [with high communalities loading with *diagnosis* (.897), *life-time axis-I psychiatric diagnostic status* (.695), and *suicide completion status* (.670) respectively)]; and finally 8.89% of the variance was explained by <u>RIN-scores</u> [with high communalities loading only with *RIN-value* (.801) and albeit weaker negative communalities loading with both sex (-.494) and post mortem index (PMI) (-.357)].

*Post-hoc* multiple comparisons of the high-order factor-analytic variables yielded group-differences in *psychiatric morbidity-&-mortality*: using Bonferroni correction, *psychiatric morbidity-&-mortality* was highest in bipolar disorder>controls (p<0.0001, **Supplementary Fig 1**); bipolar disorder>major depressive disorder (p<0.0001, **Supplementary Fig 1**), and major depressive disorder>controls (p<0.0001, **Supplementary Fig 1**). Linear regression revealed that *psychiatric morbidity-&-mortality* negatively predicted (a) the higher-order factor termed here as RIN-scores (which is an aggregate measure of RIN-values as the variable loading highest on the factor, and negative loadings of sex and PMI which loaded weakly on this factor) at (B=-2.1, t=-3.3, p=0.001) across groups; (b) fecundity at (B=-3.17, t=-2, p=0.041, *d*=1.79) across groups; and (c) age at death at (B=-8.7, t=-.27, p=0.025, *d*=2.1) only across major depressive disorder and bipolar disorder. These findings prescribed our subsequent analytical focus on *psychiatric morbidity-&-mortality* and RIN-values were included as covariates for all differential gene expression analyses. Including RIN-values as covariate was necessary especially given the negative relationship between this important measure of RNA integrity/quality (as reflected in the higher-order factor we are calling RIN-Score) and the degree of psychiatric morbidity-&-mortality across the postmortem samples.

### Differential-Expression Identified Enriched *Postmortem* Anterior Insula Gene-Expression Signatures for Mood Disorders

Differential gene-expression analyses assessed transcriptomic profiles associated with variability between cases vs controls, between high vs low *psychiatric morbidity-&-mortality,* and between *suicides vs non-suicides*. For the case-control analysis, we compared gene expression between controls>MDD and found 351 differentially expressed genes that are mainly associated with cellular homeostatic, regulation of neurogenesis, and hormonal signaling pathways (see Supplementary Table 2). We further compared controls>bipolar disorder and found 23 differentially expressed genes in mainly cellular, metabolic, hormonal signaling and neurodevelopmental pathways (see Supplementary Table 3).

For comparison of gene expression between high vs low morbidity and mortality, we binned mood disorder associated *psychiatric morbidity-&-mortality* scores into ≤ 0.82 for <u>low</u> vs ≥ 0.82 for <u>high</u> scores using a split-half method of dividing the maximum score across groups by 2 (i.e. 1.64/2) and found differentially-expressed immune and inflammatory-pathways, toll-like receptor-signaling, nuclear factor kappa-light-chain-enhancer of activated B cells (NF-kb-signaling), chemokine-signaling, and cytokine-cytokine receptor interactive pathway genes (**Fig 2-3**, and **Table 2A-F**). Specifically, within major depressive disorder and controls (i.e. major depressive disorder cases scoring high on *psychiatric morbidity-&-mortality* vs low scores and controls), we found 9 <u>under-expressed</u> inflammatory cytokine and AKT-signaling (CCL3 & CCL4); innate immunity/antigen recognition and elimination of bound antigens (IGHV2-5 and IGHV2-70); mRNA splicing and enzyme-binding (HSPA6); cellular development and homeostasis (HSPA7, PCSK5, & SERPINH1) genes; and 1 mitochondrially encoded cytochrome-C-oxidase lncRNA-pseudogene (MTCO1P12); and an <u>over-expressed</u> lncRNA-pseudogene (MTCO2P12) (**Table 2A**, **Fig 2A-D**, **Supplementary Figure 2**).

**Table 2.**
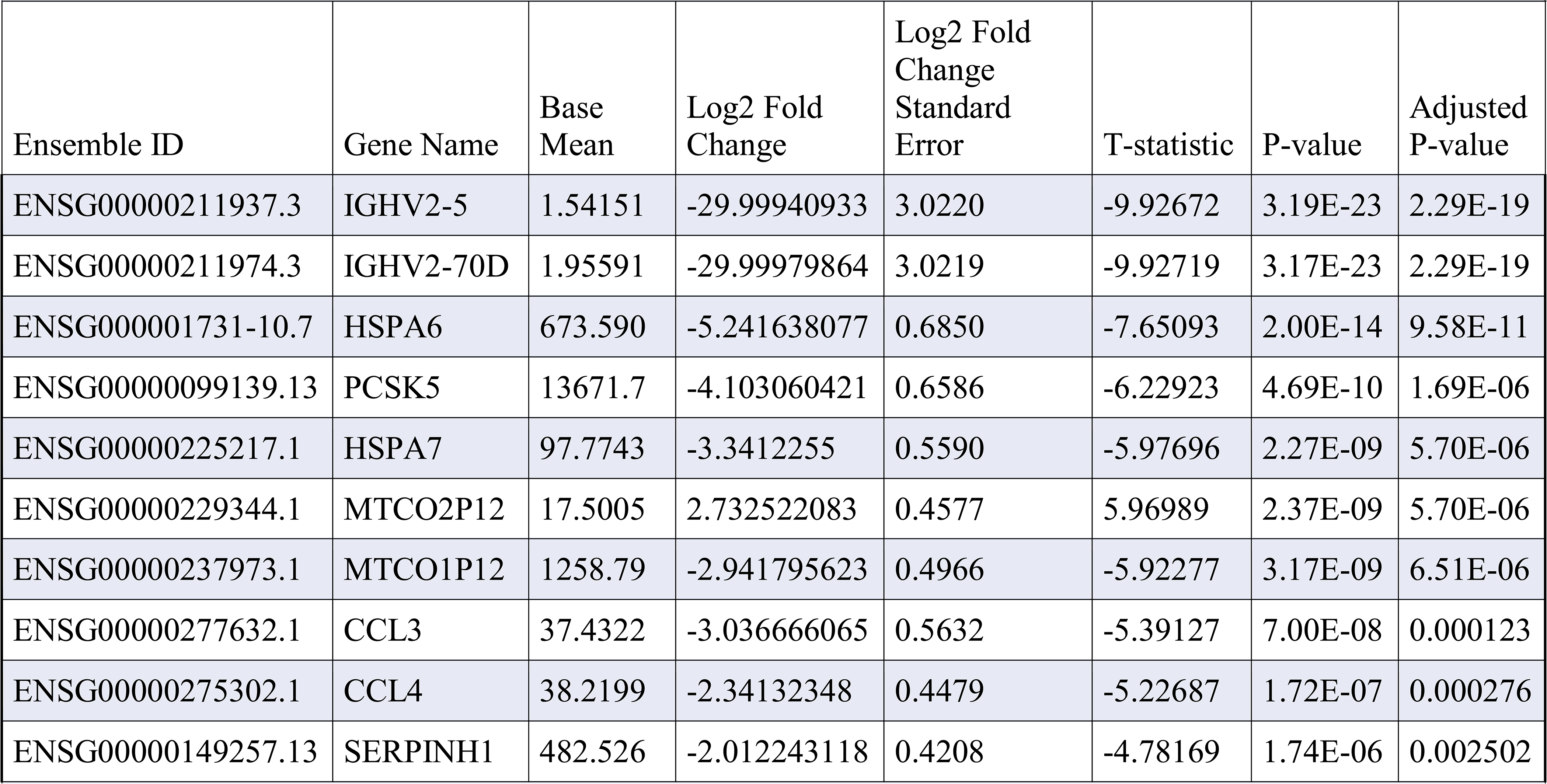
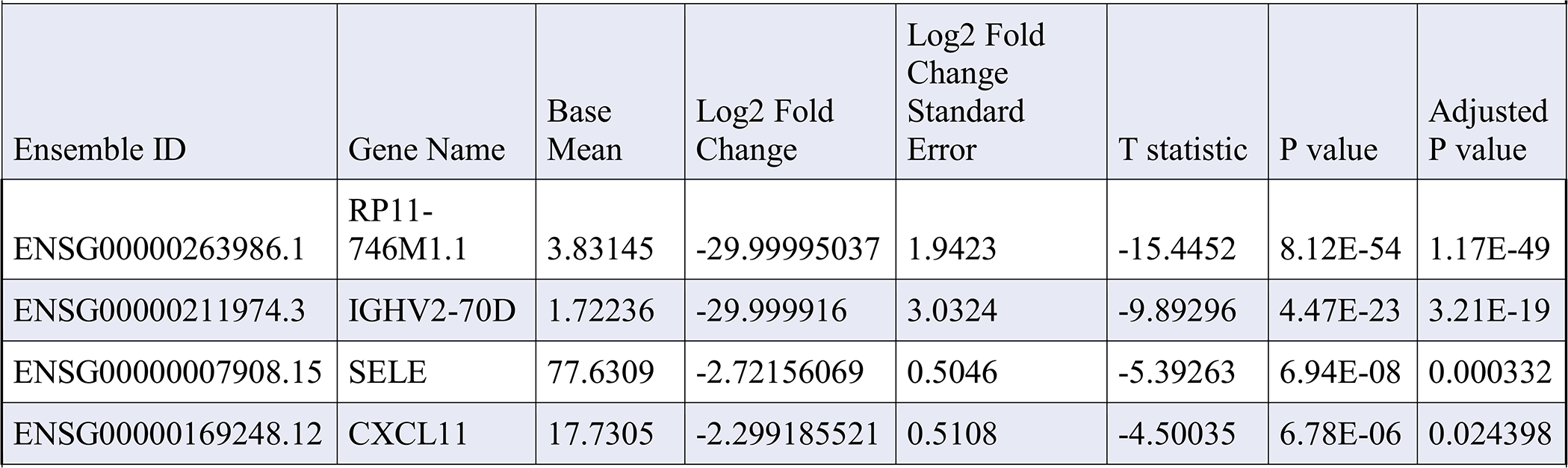
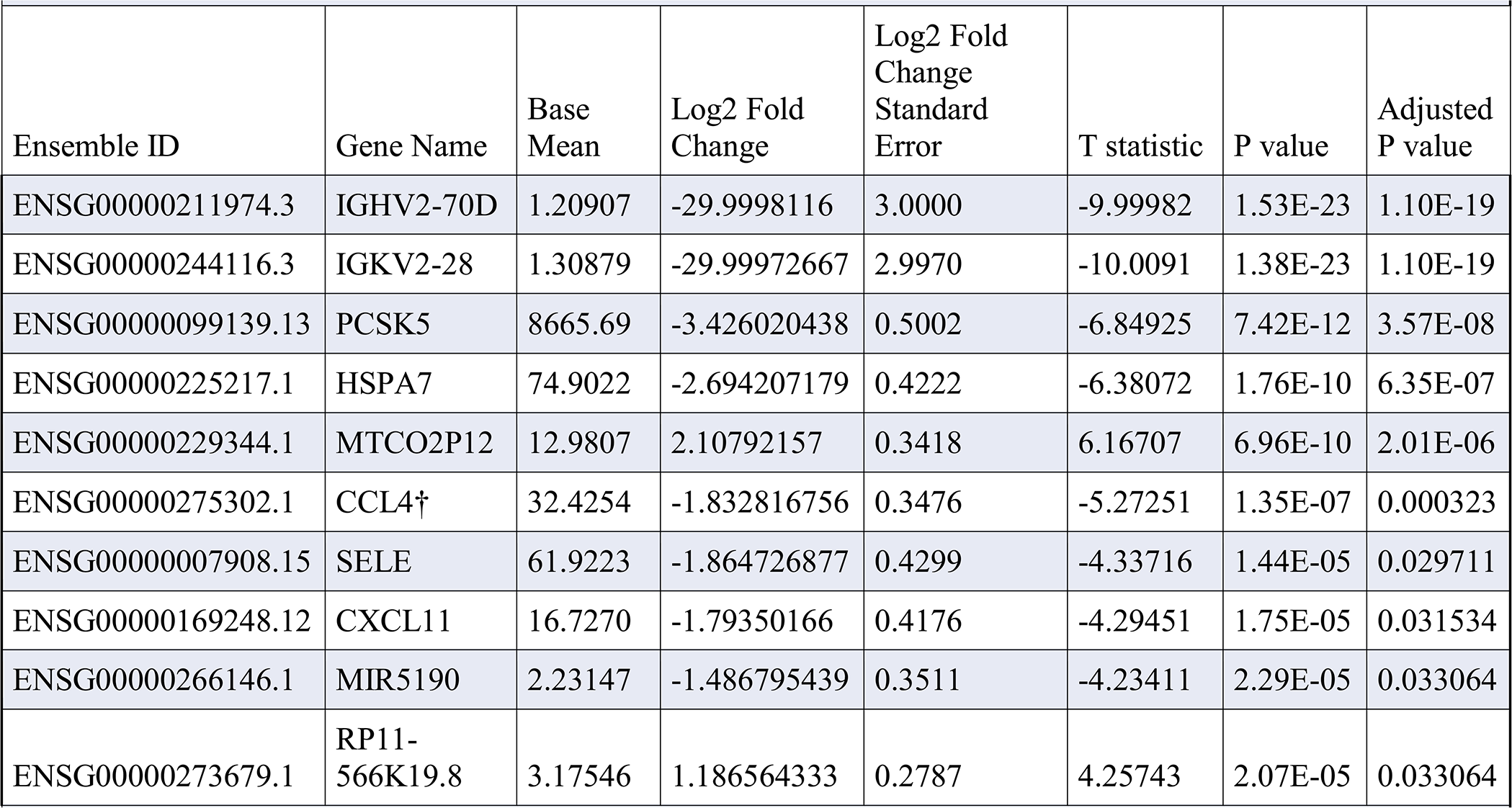
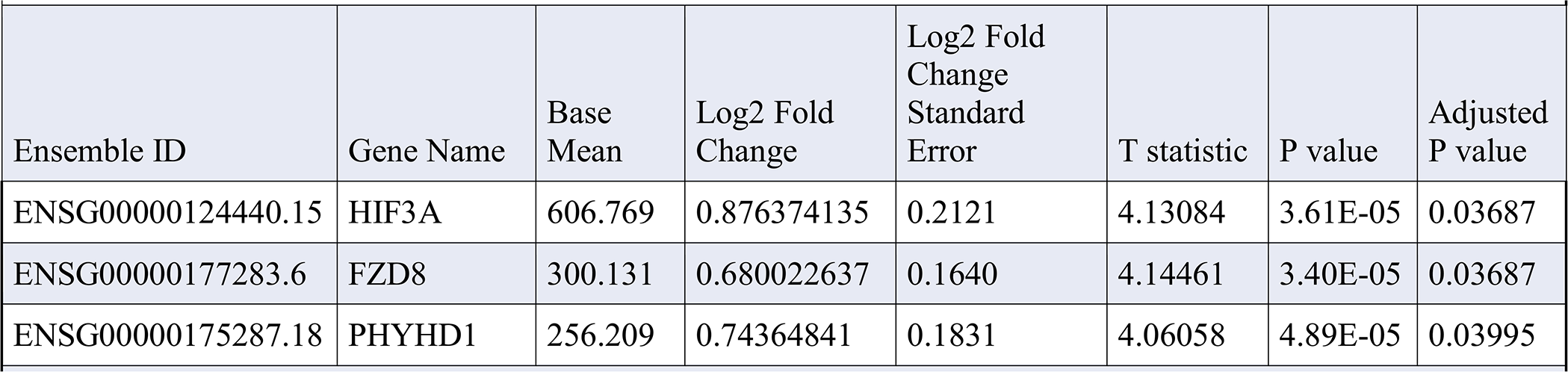
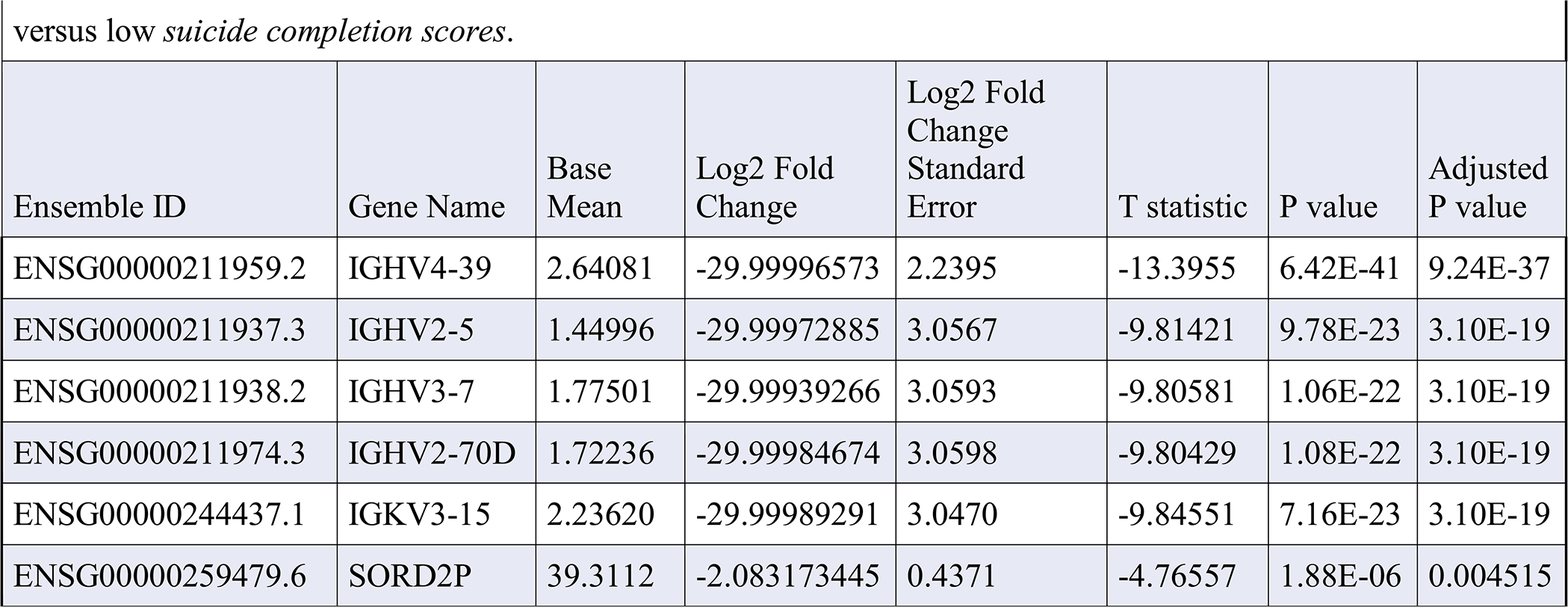
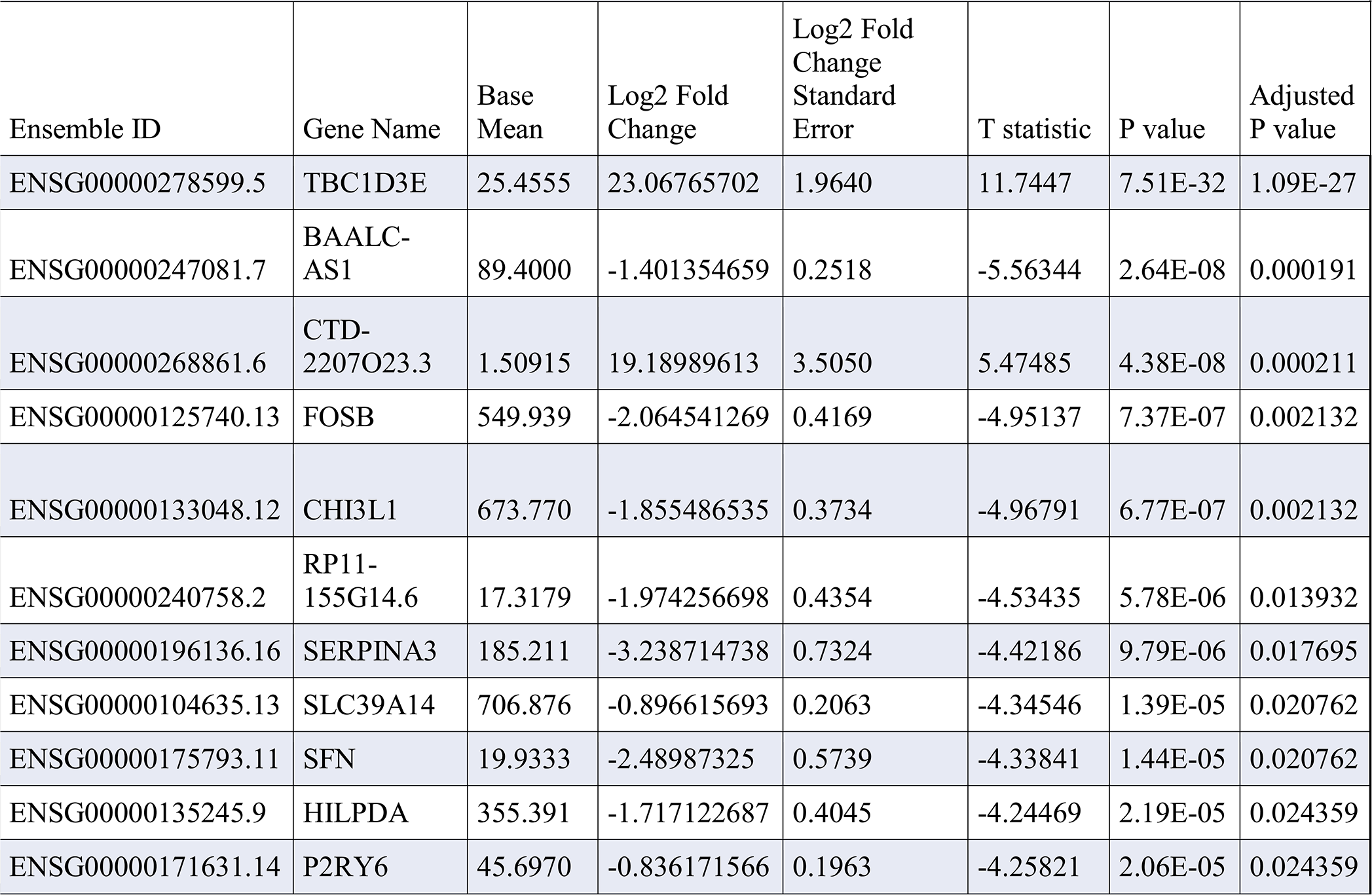
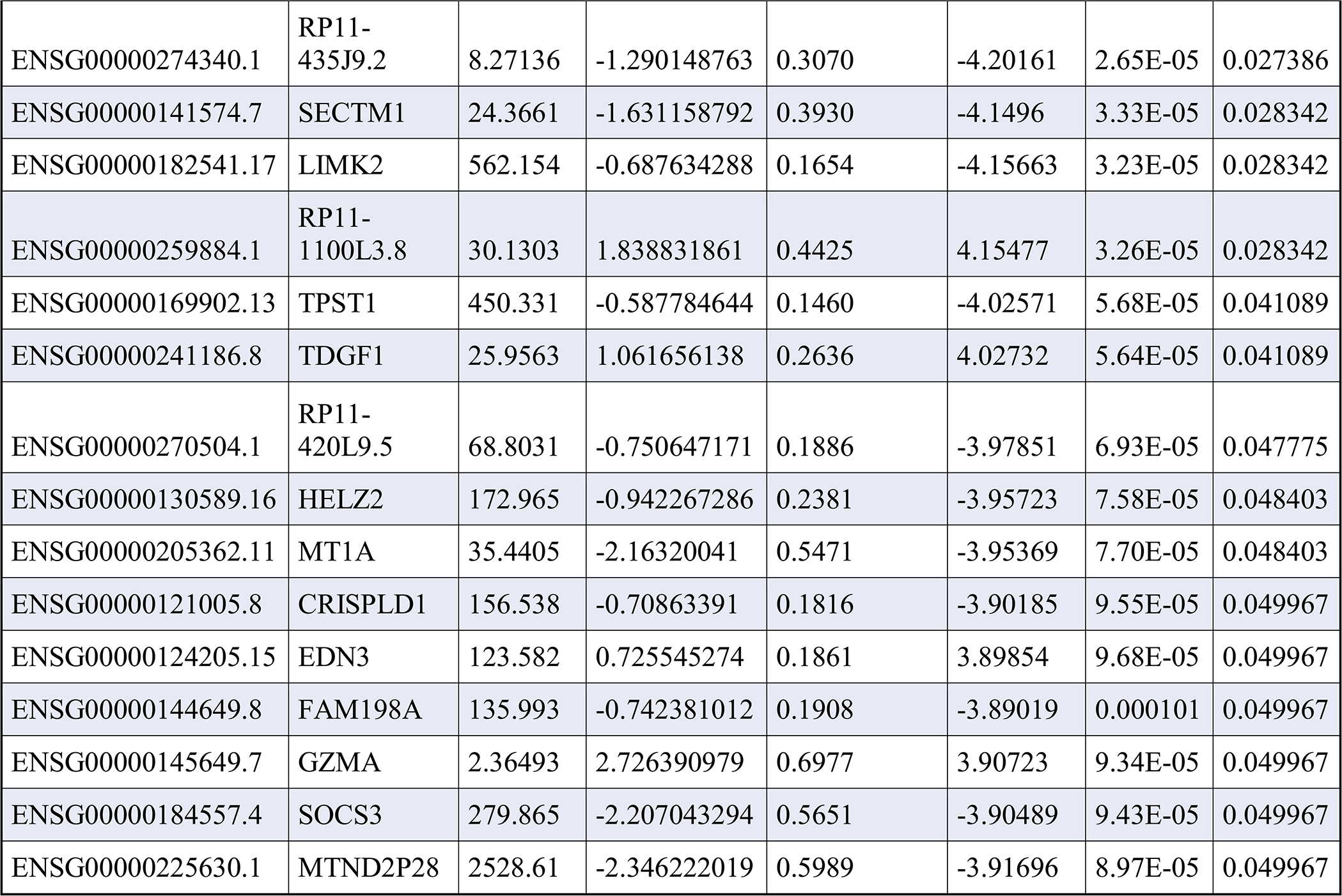
Significantly differentially expressed genes among unipolar and control samples, comparing across high versus low psychiatric morbidity and suicide-mortality scores.

**Figure 2.**
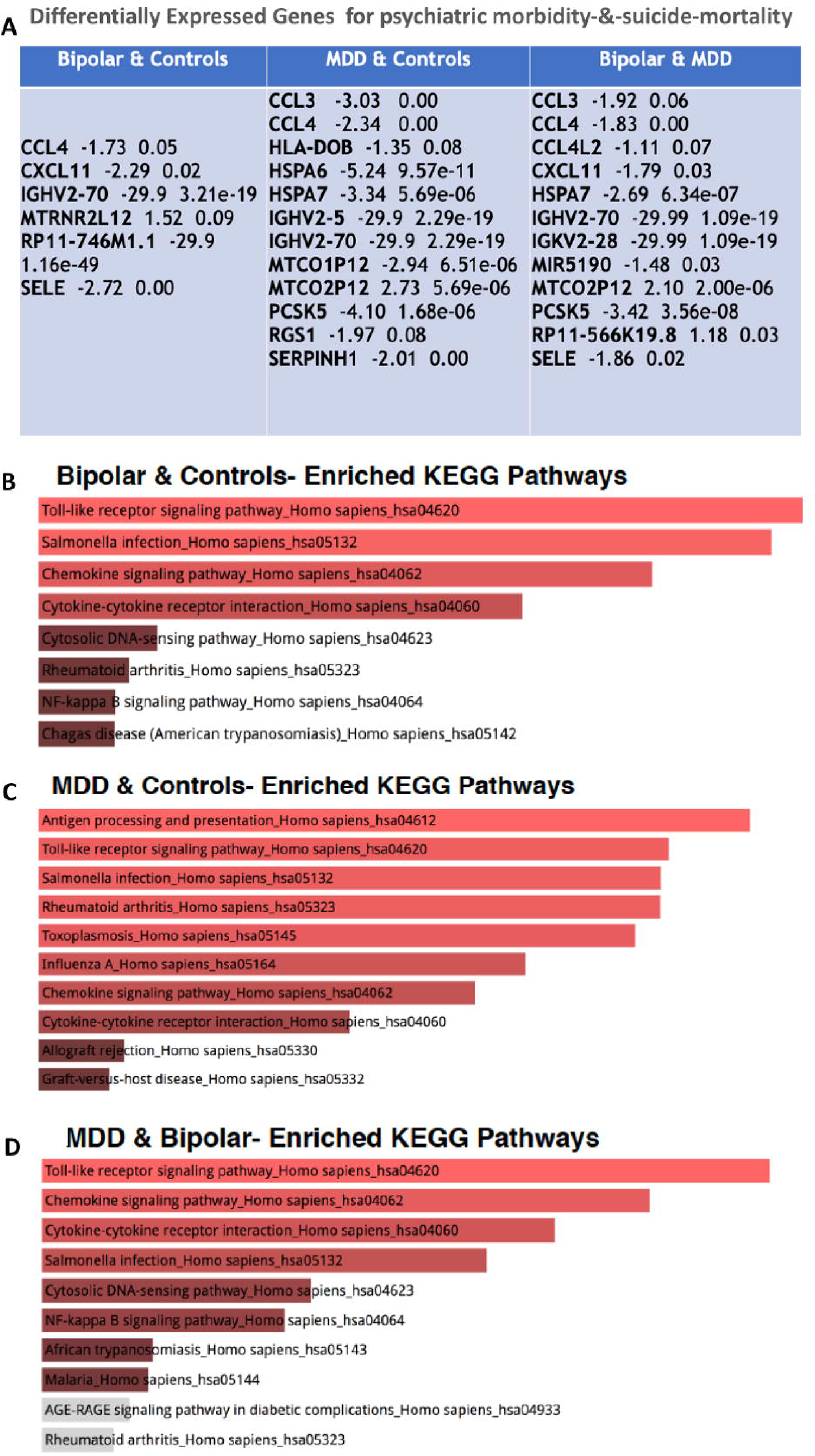
Differentially-expressed genes associated with mood disorder associated psychiatric morbidity and suicide mortality. A) shows able containing significantly differentially expressed genes identified for each comparison. The gene names are listed along with the log 2 fold change and adjusted p-value. Genes with absolute fold change >= 1.5 (here log 2 fold changes are provided) and adjusted p-values <= 0.1 were selected as significantly differentially expressed. B-D) Kyoto Encyclopedia of Genes and Genome (KEGG) pathways enriched in genes differentially-expressed between low and high suicide-mortality scores. The pathways are ranked by a combined score of p-value and rank based score. B) Bipolar and control samples only; C) major depressive disorder and control samples only; D) major depressive disorder and bipolar samples only.

A similar differential gene-expression analysis in bipolar disorder individuals with higher *psychiatric morbidity-&-mortality* scores vs those with lower scores and controls was associated with 4 <u>under-expressed</u> innate immunity (IGHV2-70); neuroprotection, neurodevelopment and CNS-diseases including major depressive disorder (CXCL11, RP11-7461.1, & SELE) pathway genes (**Table 2B**, **Fig 2A-D**, **Supplementary Figure 3**). Analysis of **high** *psychiatric morbidity-&-suicide-mortality* scores in the pooled major depressive disorder and bipolar disorder (i.e. mood disorders) vs **low** *psychiatric morbidity-&-mortality* scores in the pooled mood disorders and controls yielded 10 <u>under-expressed</u> inflammatory cytokine and AKT-signaling (CCL4); innate immunity (IGHV2-70 & IGKV2-28); cell-neurodevelopment and CNS diseases (CXCL11, SELE, PCSK5, & HSPA7); and transcriptional regulatory-RNA (MIR5190); but also 2 <u>over-expressed</u> innate immunity (RP11-566K19.8) pathway genes; and a lncRNA-pseudogene (MTCO2P12) (**Table 2-C**, **Fig 2A-D**, **Supplementary Figure 4**).

We then examined gene-expression profiles related to suicide by comparing suicide completers vs non-suicide completers with major depressive disorder and controls (i.e. comparing gene-expression in major depressive disorder suicides vs major depressive disorder non-suicide and controls), and found 3 <u>over-expressed</u> WNT-signaling (FZD8); transcriptional regulation of adaptive responses to oxygen tension/hypoxia, DNA-binding transcriptional activity/co-activation (HIF3A); and dioxygenase activity (PHYHD) pathway genes (**Table 2D**). Comparing gene expression profiles for suicide completion in bipolar disorder-suicides vs non-suicidal bipolar disorder and controls, we found 5 <u>under-expressed</u> innate immunity (IGHV2-5, IGHV2-70, IGHV3-7, IGHV4-39, & IGHV3-15); and CNS-disease (SORD2P) pathway genes (**Table 2E**).

Assessing gene-expression profiles associated with suicide-completion in the pooled major depressive disorder and bipolar disorder completed-suicides vs major depressive disorder and bipolar disorder non-suicidal deaths yielded 21 <u>under-expressed</u> innate immune and inflammatory-cytokine (CRISPLD1, CHI3L1, P2RY6, & SECTM1); protein-protein interaction regulatory (MT1A, HILPDA, HELZ2, FOSB, FAM198A, SOCS3, & TPST1); neurodegeneration (RP11-155G14.6, SLC39A14, & SERPINA3); cellular-neurodevelopmental and transcriptional (LIMK2, SFN, & EDN3) pathway genes (**Table 2F**); as well as uncharacterized genes/pseudogenes (MTND2P28, BAALC-AS1, RP11.420L9.5, & RP11.435J9.2). We also found 5 <u>over-expressed</u> inflammatory (RP11.1100L3.8); intracellular protein transport (TBC1D3E); cell fate and apoptosis regulation (GZMA); and transcriptional, embryonic/forebrain cell development and defect (CTD-2207O23.3, & TDGF1) pathway genes (**Table 1F**, **Fig 2D**).

### WGCNA Identified co-regulated gene modules for *Postmortem* Anterior Insula Gene-Expression Signatures for Mood Disorders

WGCNA characterized the potential co-regulated gene-modules involved in mood disorder *morbidity and suicide-mortality*. After correcting for covariates and filtering out low-mean genes, we found 2 dominant co-expression modules (coded as **blue** and black **Figs. 3-4; Supplementary Figure 5**) strongly associated with *psychiatric morbidity-&-mortality* variables.

Using the 30 most highly connected (i.e. hub genes) in these two modules, the Blue module showed enrichment in genetic pathways involved in fundamental cellular signaling processes such as ATPase activity, cAMP signaling, sodium and calcium channel functions, transcriptional regulation, and neuronal development/proliferation and differentiation (**Figs. 3A-C; Supplementary Figure 5)**. The Black module showed enrichment in genetic pathways involved in innate immune functions, cellular homeostasis and metabolic regulations, dopamine DARPP32 feedback onto cAMP functions in the regulation of dopaminergic synapse, glial cell development, addiction and major depressive disorder risk (**Fig. 3A-C** and **Fig. 4A-B**; **Supplementary Figure 5**).

**Figure 3.**
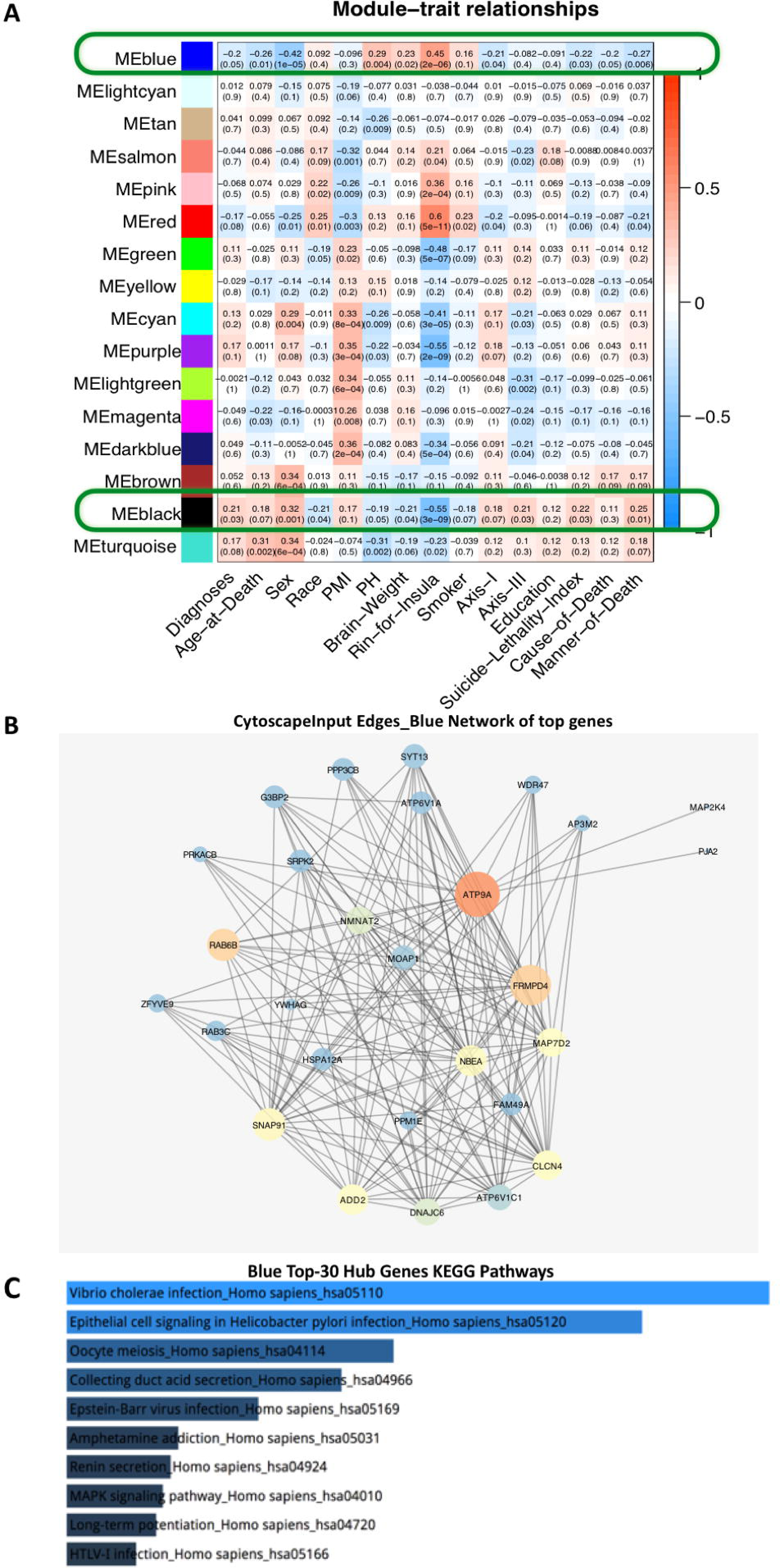
WGCNA Identified clinical and demographically-associated gene co-expression modules. A). Heatmap showing the relation of the WGCNA identified Blue and Black modules of interest (zoomed in correlation values in Blue and Black lines) with sample traits shown in the x-axis. Color scale (red-blue) represents the strength of the correlation between the module and the trait. Correlation values (R^∧^2) followed by corresponding p-values (in parenthesis) are listed within the heatmap. Other modules shown in A have greater negative association with biological covariates (i.e. RIN, sex and race, & technical covariates such as pH). B) Hub Genes from WGCNA Blue Module: Genes with the highest connectivity (larger circles) to other nodes within the blue module (smaller circles) within the co-activated network of genes were identified as hub genes. Gene-Gene network constructed using the hub genes with the size of nodes scaled by degree and color of nodes scaled by betweenness centrality (darker colors for lower values). C) Enriched KEGG pathways for the 30 hub genes for the Blue module. The pathways are ranked by a combined score of p-value and rank based score. WGCNA identifies multiple modules of genes that are co-expressed. The module’s eigengene which is representative of the expression of that module is correlated to different metadata variables to identify modules of interest. We correlated our modules to different metadata variables of postmortem sample characteristics and selected the two modules which showed significant correlation to variables of interest such as diagnoses, suicide, and the lethality of suicide methods.

**Figure 4.**
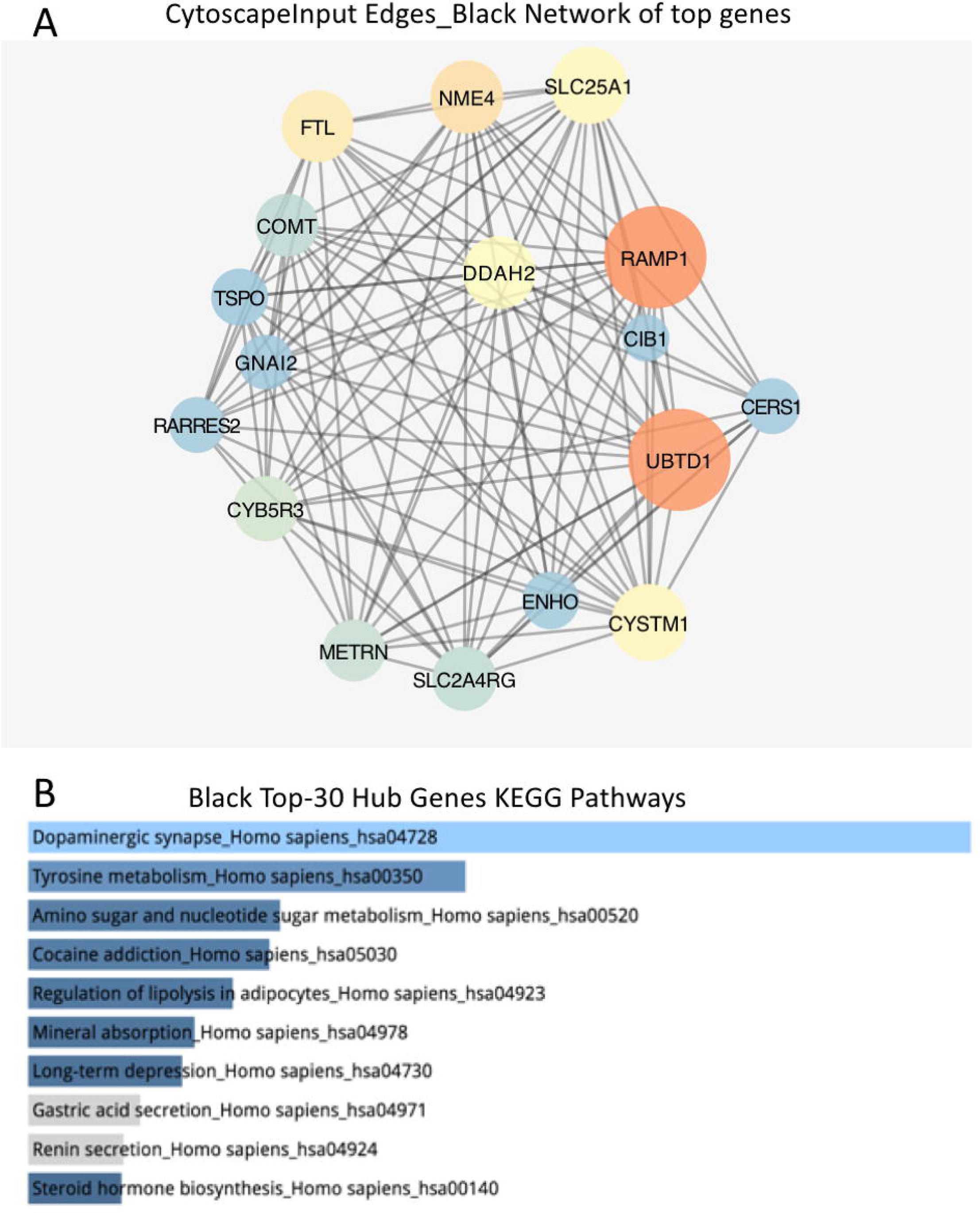
WGCNA Identified clinical and demographically-associated gene-module – Black Module. A) Gene-Gene network constructed using the hub genes with the size of nodes scaled by degree and color of nodes scaled by betweenness centrality (darker colors for lower values. B) Enriched KEGG pathways for the 30 hub genes for the black module. The pathways are ranked by a combined score of p-value and rank based score.

## DISCUSSION

Using a neuroimaging meta-analysis to refine a structurally reduced AIC region of interest in major depression and bipolar disorder, we identified a *postmortem* across-diagnostic mood disorder linked psychiatric morbidity-&-mortality associated gene-expression signature within this neuroimaging meta-analysis identified reduced AIC gray matter signature. Given this AIC sub-region’s documented role in regulating affective and physical pain/distress, general bodily homeostatic and interoceptive salience, our convergent structural neurobiological and functional gene-expression findings of (a) a reduced right AIC gray matter signature in living mood disorder patients, coupled with (b) a preponderance of under-expressed, but also albeit to a lesser extent, over-expressed gene-expression signatures within the identified AIC-locale in mood disorder *postmortem* brains, identified an anatomically-precise functional gene-expression basis for mood pathologies. Of note, we found a negative relationship between measures of AIC tissue RNA integrity/RIN and mood disorder morbidity, which could have implications for our results even though we corrected for RIN as a covariate. Whether our observed negative correlations between RIN measures of AIC tissue and mood disorder morbidity and suicide mortality factors is regionally dependent needs to be assessed in future studies.

In light of the right AIC’s role in sensing emotional/physical pain associated with social isolation and disconnection (Eisenberg 2015), these results provides a potential gene-regulatory windows into the neuropathological ramifications for increased mood disorder associated psychiatric morbidity (O’Connor & Nock 2014; Nock et al. 2018), which could compound suicidal outcomes (Nock et al. 2018). The right AIC’s role in coding affective salience and psychological pain (Schneidman 1998), and the possible collective modulatory impact of these states on maladaptive impulses such as the urge to escape unbearable misery via suicide, provides a putative anatomical framework for mood disorder associated psychiatric morbidity-&-mortality (Schneidman 1998; Craig 2009; Slavich et al. 2010; Mee et al. 2011; Hatton et al. 2012; McGrath et al. 2013; Wager et al. 2013; Eisenberg 2015; Goodkind et al. 2015; Koechlin 2016; Wise et al. 2017; Khalsa et al. 2018). The identified association between mood and associated psychiatric morbidity-&-mortality scores with fecundity, and with predominant under-expressed gene-expressive functions suggests an evolutionary significance of the current results. For instance, genetic abnormalities governing aberrant AIC-mediated social deficits (Jabbi et al. 2012) and severe mood disorders could attenuate reproductive prowess (Mullins et al. 2017).

Within the identified reduced AIC signature, our differential gene-expression analyses identified predominant down-regulations in innate immune functions, inflammation pathways, and AKT-signaling related to mood-associated psychiatric morbidity and suicidal-mortality.

Further differential-expressions involving cellular-homeostatic, neurodevelopmental, and transcriptional pathways in the AIC were found to be associated with psychiatric morbidity and suicidal-mortality in less directionally-specific (i.e. up/down-regulatory) patterns. While the cellular-origins of our findings of predominantly downregulated immune-related and neurodevelopmental gene-expression changes cannot be specified with our bulk tissue RNA-seq methods, astrocyte-derived cytokine functions have been documented to induce synapse engulfment/elimination and thereby influence synaptic pruning (Vianchtien et al. 2018; Bennett & Molofsky 2019). Furthermore, immune pathway mediation of mood dysfunctions and related psychiatric diagnoses are proposed to be likely multifaceted as part of the brain’s immune-related response repertoire such as toll-like receptor signaling, can be influenced by both ***a***) pathogen-associated molecular patterns (Kawai & Akira 2007) and ***b***) danger-associated molecular patterns (Klune et al. 2008; Piccinini et al. 2010). This perspective on the multifaceted nature of brain immune signaling in relation to behavioral dysfunctions like mood disorders deserves further analysis especially in light of *1*) the strong link between endogenous danger-associated immune/inflammatory cellular functions in promoting homeostasis (Klune et al. 2008) and *2*) the potential impacts of environmental stress experiences on endogenous cellular stress as well as inflammatory responses (Slavich et al. 2010).

Existing findings of both over-expressed immune-related signals (Pandey GN 2017), and our current imaging-guided AIC postmortem findings that clearly replicated previous results of predominantly under-expressed immune/inflammatory function (e.g. chemokine ligand 2 CCL4) and genes/pseudogenes implicated in regulatory cellular functions (e.g. a serpin peptidase inhibitor & HSPA7) in postmortem dorsolateral prefrontal cortex BA9 brains of mood disorder individuals with and without suicide completion (Pantazatos et al. 2017), needs to be better contextualized. For instance whether these differences in the directionality of regulatory patterns of gene-expression findings (i.e. over-expressed vs under-expressed immune expressive genes) in mood disorder postmortem studies is somewhat related to methodological differences in in terms of targeted micro-RNA assays vs whole transcriptome sequencing approaches, or differences in sample sizes, sample selection criteria, or qualitative postmortem material differences between studies, needs to be examined more carefully.

While we cannot fully disentangle medication effects from diagnosis effects in our current preliminary analysis, it is worth noting that our observations suggest that diagnostic effects relating to both our suicide related and non-suicide related gene expression results are likely manifold, beyond mood disorders. This is relevant especially in light of our sample characteristics showing that suicide completion is collinear with mood disorder diagnostic severity, and that suicide completers are about 80% of the included cases. Our findings of consistently similar gene expression patterns across the various morbidity & mortality comparisons further support the thesis that suicide completion and mood disorder morbidity are likely more inter-related at the biological level than previously appreciated. Together, our current findings are consistent with data on the role of immune dysfunctions in CNS diseases (Oquendo et al. 2014; Wohleb et al. 2016; Pantazatos et al. 2017; Butovsky et al. 2018), and inflammasome functional prediction of major depressive disorder treatment outcomes (Syed et al. 2018). The results further lend credence to the hypothesis that neurodevelopmental and transcription-factor genes are critical mediators of complex adaptive brain functions (Changeaux et al. 2017); especially within the context of the AIC’s integration of affective and physiological feeling states (Craig 2009; Slavich et al. 2010; Kurth et al. 2010; Eisenberg 2015; Khalsa et al. 2018) ‘including homeostatic maintenances in sickness and health’ (Craig 2009; Khalsa et al. 2018), that are likely not entirely independent of both pathogen-associated molecular patterns (Kawai & Akira 2007) and danger-associated molecular patterns (Klune et al. 2008; Piccinini et al. 2010) known to induce brain immune signaling.

At the systems level, the toll-like receptor (TLR) pathway genes found to be under-expressed here are documented to recognize conserved motifs in microorganisms (Akira 2003) and stimulation of TLRs are shown to mediate acute-immune defense and cytokine production/release (Perkins 2007). First, it is noteworthy that the immunoglobulin heavy chain variable region 2 (IGHV2) pathway genes have shown to be more than 20-fold differentially expressed in donors with low vs. high mood disorder morbidity (Axis-I diagnostic load) and suicide mortality in the current study. This observation of consistent under-expression of IGHV2 genes in bipolar disorder vs controls, MDD vs controls, and the pooled mood disorders vs controls suggests a possible role for the IGHV2 pathway in mood disorder morbidity and mortality, at least in the current anterior postmortem sample. Furthermore, the observed under-expression/suppression of innate immune pathway genes in higher morbidity and suicide mortality afflicted mood disorder donors point to this pathway’s possible transcriptomic dysregulatory properties in depressive illness. This is especially in light of evidence that depression was the most recent episode in 21 of the 37 bipolar donors coupled with data that 26 of the 30 MDD donors have a lifetime history of recurrent depressive episodes in the current samples. To our knowledge, there are no previous reports implicating the IGHV2 pathway gene expression or genomics in mood or depression pathogenesis. However, examining the role of IGHV2 and related innate immune pathways in studies including larger mood disorder samples including more than one brain region, will help confirm their possible fundamental roles in mood illness transcriptomics.

Second, our identified under-expressed NF-κB pathway genes are implicated in controlling DNA transcription, cytokine production and cell survival (Meffert et al. 2003), and are essential for cellular-immune response to infection, stress-related shocks (Van Amerongen et al. 2009), and synaptic plasticity and memory (Meffert et al. 2003; Van Amerongen et al. 2009). Of interest, NF-kB pathway genes are also identified as a critical mediator of stress-impaired neurogenesis and depressive-like behavior caused by exposure to chronic stress in mice (Koo et al. 2010). Furthermore, recent studies identified an NF-kB pathway involvement in the pathophysiology of depressive illness, especially relating to neurogenesis, synaptic transmission and plasticity (Caviedes et al. 2017). NF-kB related mechanisms have also been shown to have antidepressive therapeutic effects in general (Wu et al. 2018) and in inflammatory disease comorbidity with depression-like symptoms in mice (Su et al. 2017).

Third, the identified under-expressed chemokine-signaling pathways govern critical spatiotemporal cell-positioning during developmental coordination and translational guidance of cell-locomotion and migration (Turner et al. 2014). Fourth, the identified under-expressed cytokines are implicated in cell-specific innate and adaptive inflammatory host defenses, cellular-development, cell-death, angiogenesis, and maintenance of cellular homeostasis (Syed et al. 2018). Conversely, the Wnt-β-catenin signaling pathway found to be over-expressed in major depressive disorder suicides is an evolutionarily conserved inter-cellular communication system that mediates stem cell renewal, cell-proliferation and differentiation during embryogenesis and adulthood (Meffert et al. 2003). Taken together, our observations of convergent under-expressed

TLRs, NF-κB, chemokine, and cytokine-cytokine interactive pathways transcriptomic signatures for psychiatric morbidity-&-mortality; and suicide-mortality-specific over-expressed Wnt signaling pathway, suggests possible dysregulatory mechanisms for aberrant cellular processes very early in development. These processes may negatively shape adaptive immune, inflammasome and chemokine-cytokine responses to adverse socio-emotional and environmental distress, with a prolonged experience of these adverse circumstances likely leading to compromised AIC anatomical and physiological integrity, and associated maladaptive rupture in regulatory mood states.

Of relevance, our AIC postmortem WGCNA results which reflects a global perspective whole transcriptome gene expression by identifying co-expressed gene-modules for *a*) lifetime mood disorder-diagnoses, *b*) lifetime Axis-I diagnoses, and *c*) suicide-completion status as well as the lethality of the committed suicide methods in genetic pathways involved in fundamental cellular signaling processes like transcriptional regulation, ATPase and cAMP signaling, sodium and calcium channel functions, neuronal developmental processes (Blue module); as well as capturing co-expressed gene-modules *a*) lifetime mood disorder-diagnoses, *b*) lifetime Axis-I diagnoses, and *c*) suicide-completion in genetic pathways involved in innate immune functions, cellular homeostasis and metabolic regulations, regulation of dopaminergic synapse, glial cell development, addiction and major depressive disorder risks. While the regional specificity of these findings needs further analysis, the WGCNA results recapitulate the differential gene expression findings and further underscore the complex interactive gene-functions implicated in brain mediation of mood disorder morbidity and related suicidal risk tendencies.

In line with our differential gene expression and WGCNA findings of neurodevelopmental, immune, inflammatory, transcriptional, ATPase and cellular and hormonal signaling pathway alterations in mood disorder morbidity and suicides, previous gene expression studies of the postmortem prefrontal brain system of completed suicides found altered ATPase signaling in cases (Pantazatos et al. 2017). Further studies of suicide victims with and without major depression found extensively altered limbic and hippocampal gene expression where processes linked to major depression were associated with abnormal transcription and cellular metabolic processes (Sequeira et al. 2007). At the synaptic level, altered gene expression related to intracellular signaling in GABAergic (Sequeira et al. 2007; Sequeira et al. 2009) and glutamatergic transmission pathways (Sequeira et al. 2009) have also been reported. Gene set enrichment analysis and expression pattern exploration in postmortem brains across the lifespan in bipolar disorder has recently implicate enrichments of neurodevelopment processes in bipolar illness (Mühleisen et al. 2018). Combining differential expression and WGCNA, a recent postmortem study identified cross-species major rearrangement of transcriptional patterns in the ERK signaling and pyramidal neuron excitability in major depressive illness, with limited overlap in males and females in both depressed humans and chronic stressed mice (Labonté et al 2017). Taken together, substantive evidence of ATPase, cellular processes and signaling, and neurodevelopmental pathway gene expression abnormalities found in limbic and prefrontal brain regions in previous studies are in line with our current findings of AIC postmortem gene expression profiles associated with mood disorder morbidity and mortality. Future studies including multiple brain regions and sex-specific analyses will be needed to provide in-depth mechanistic understanding of the underlying transcriptional landscapes associated with diagnostic and sex-specific aspects of mood illnesses and related suicide mortality.

Unlike cardiovascular disease and cancer research, where pathobiological measures are causally linked to disease morbidity and end-point mortality, the causal neurobiological root causes of mental illnesses and their associated mortality endpoints such as suicides are unknown, limiting measurable biological predictability of suicidal-mortality. Our findings of convergent structural neurobiologically defined functional gene-expression signatures for mood disorder associated psychiatric morbidity-&-mortality across major depressive disorder and bipolar disorder supports shared heritable neurogenetic pathologies underlying comorbid neuropsychiatric symptoms (Anttila et al. 2018; Gandal et al. 2018). While the cell-type specific aberrations and their relationship with differential gene expression profiles needs to be studied to better understand the molecular mechanisms underlying abnormal neuroanatomical signatures for mood symptoms, especially in-terms of diagnostic specificity between major depressive disorder and bipolar disorder, our results represent a step towards developing brain region-specific functional gene-expression blueprints for therapeutic targeting of broad/specified molecular pathways. Furthermore, while we are not able to disentangle medication effects in our current findings, the effects of medication on gross neuroanatomical measures and gene-expression profiles also needs to be assessed in future studies. Our analysis of suicide vs non-suicide mood disorders (excluding controls) did compare similarly medicated diagnostic cohorts and as such, our preliminary results showing similarities in gene expression profiles between suicide comparisons with and without controls suggests that our findings overall are not attributable to medication effects alone. In sum, our findings bridging convergent neuroanatomical and gene-expression signatures for measures of the degree of comorbid psychiatric symptoms in mood disorders and suicides, represents a framework for discoveries of novel biomarkers for brain diseases.

## Author Contributions

MJ conceived and designed the studies, and acquired postmortem material from the NIMH HBCC. MJ, SBE, DA and HH performed the experiments and analyzed the data and results. MJ drafted the manuscript and DA, SBE, SMS, CBN, and HH contributed critically to the interpretation of the findings and writing the paper.

## Acknowledgements

The NIMH Human Brain Collection Core provided RNA-samples for all 100 *postmortem* data and we thank the NIMH and Drs. Barbara Lipska, Stefano Marenco, Pavan Auluck and HBCC colleagues for providing the studied samples. We thank Wade Weber of Dell Medical School Psychiatry Department, UT Austin for assistance in preparing the manuscript, Dr. Mark Bond of Dell Medical School Psychiatry Department, UT Austin for statistical reviews, Nicole Elmer of the Biomedical research support for help with gene-expression result figures, and Jessica Podnar and several GSAF colleagues for RNA-seq support. This work was supported by the Dell Medical School, UT Austin startup funds and Mulva Neuroscience funds for MJabbi, and HHofmann is supported by NSF-DEB 1638861 and NSF-IOS 1326187.

## Potential Conflicts of Interest

Mbemba Jabbi, none. Dhivya Arasappan, none. Simon Eickhoff none.

Hans Hofmann, none.

Stephen Strakowski: chairs DSMBs for Sunovion as a Consultant, and has research grants with Janssen pharmaceuticals and Alkermes.

Charles Nemeroff: 1) Consulting (*last three years*) for Xhale, Takeda, Taisho Pharmaceutical Inc., Bracket (Clintara), Fortress Biotech, Sunovion Pharmaceuticals Inc., Sumitomo Dainippon Pharma, Janssen Research & Development LLC, Magstim, Inc., Navitor Pharmaceuticals, Inc., TC MSO, Inc., Intra-Cellular Therapies, Inc; 2) Stockholder for Xhale, Celgene, Seattle Genetics, Abbvie, OPKO Health, Inc., Antares, BI Gen Holdings, Inc., Corcept Therapeutics Pharmaceuticals Company, TC MSO, Inc., Trends in Pharma Development, LLC; 3) Scientific Advisory Board member for American Foundation for Suicide Prevention (AFSP), Brain and Behavior Research Foundation (BBRF), Xhale, Anxiety Disorders Association of America (ADAA), Skyland Trail, Bracket (Clintara), Laureate Institute for Brain Research (LIBR), Inc; 4) Board of Directors member for AFSP, Gratitude America, ADAA; 5) Income sources or equity of $10,000 or more from American Psychiatric Publishing, Xhale, Bracket (Clintara), CME Outfitters, Takeda, Intra-Cellular Therapies, Inc., Magstim, EMA Wellness; 6) Patents: Method and devices for transdermal delivery of lithium (US 6,375,990B1); Method of assessing antidepressant drug therapy via transport inhibition of monoamine neurotransmitters by ex vivo assay (US 7,148,027B2); Compounds, Compositions, Methods of Synthesis, and Methods of Treatment (CRF Receptor Binding Ligand) (US 8,551, 996 B2).

## Notes

#### Summary of Updates

Additional comparisons to include case-control differences in gene expression and clarifications of previous analyses have been included in this revision.

