## Supplementary Materials for "Neuro-transcriptomic signatures for mood disorder morbidity and suicide mortality"

#### RESULTS

##### Postmortem Group Differences and Factor Analysis

Demographic variables did not differ among groups, except for race: major depressive disorder and bipolar disorder groups had more Caucasian donors whereas healthy subjects had more African American donors (p<0.0001, F=12). Covarying for race in subsequent ANOVAs, Axis-I load (p<0.0001, F=30) (Fig 2A) and suicide-completion (p<0.0001; F=39.7) were higher in major depressive disorder and bipolar disorder donors than controls (Fig 2B). Using Bonferroni correction, post-hoc pairwise-comparisons of the original postmortem variable yielded group differences in Axis-I-load in major depressive disorder>controls (p<0.0001); bipolar disorder>controls (p<0.0001); and bipolar disorder>major depressive disorder (p = 0.004); while 14 suicide-completion differed in major depressive disorder>controls (p<0.0001) and in bipolar disorder>controls (p<0.0001), but not between major depressive disorder and bipolar disorder. Linear regression analysis showed that Axis-I-load predicted suicide-completion (B=.195, t=5.2, p<0.0001, d=1.408), but not age at death.

**Supplementary Figure 1.**

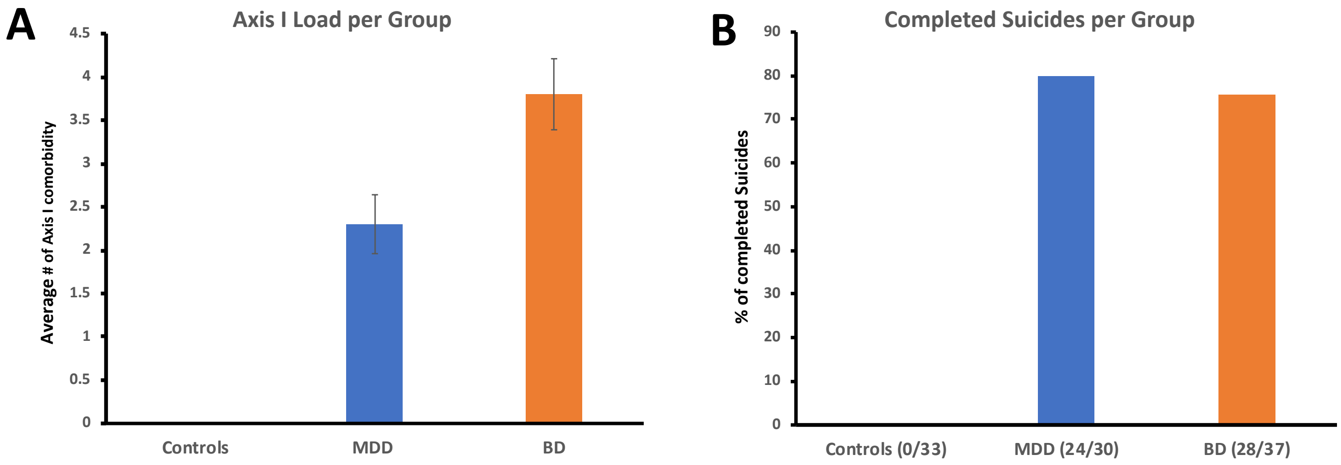


**Supplementary Figure 1. Postmortem psychiatric morbidity as shown in Axis-I comorbidity load & percentage of suicide completion across diagnostic groups.** A) Shows the mean lifetime Axis-I diagnostic load across the major depressive disorder, bipolar disorder and control donors; with the bipolar disorder group showing the highest comorbid Axis-I diagnostic load relative to major depressive disorder and controls. B) Suicide completion as shown in percentage of deaths by suicide in the major depressive disorder (24 suicides out of 30 deaths) and bipolar disorder (28 suicides out of 37 deaths) groups. Suicide was an exclusion criteria for controls. Error bar for A represents SEM.

### Supplementary Figure 2

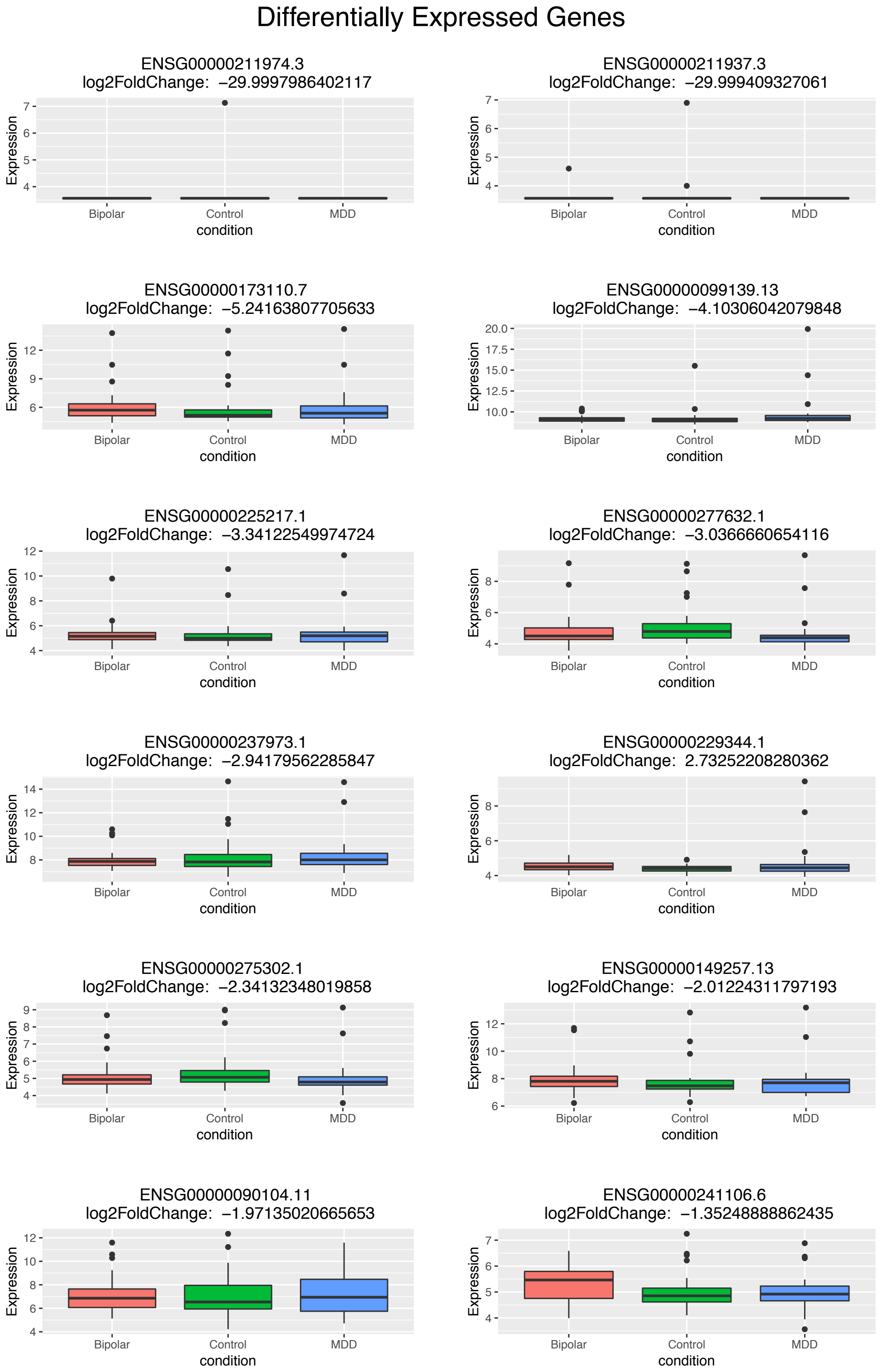


**Supplementary Figure 2**. Differential Gene Expression associated with psychiatric morbidity-&-suicide-mortality: high MDD vs low MDD & Controls. Boxplots illustrates group-specific expression mean expression values for each of the differentially expressed genes shown in Table 2A and Figure 2A. Error bars denote log-fold-change standard errors.

#### Supplementary Figure 3

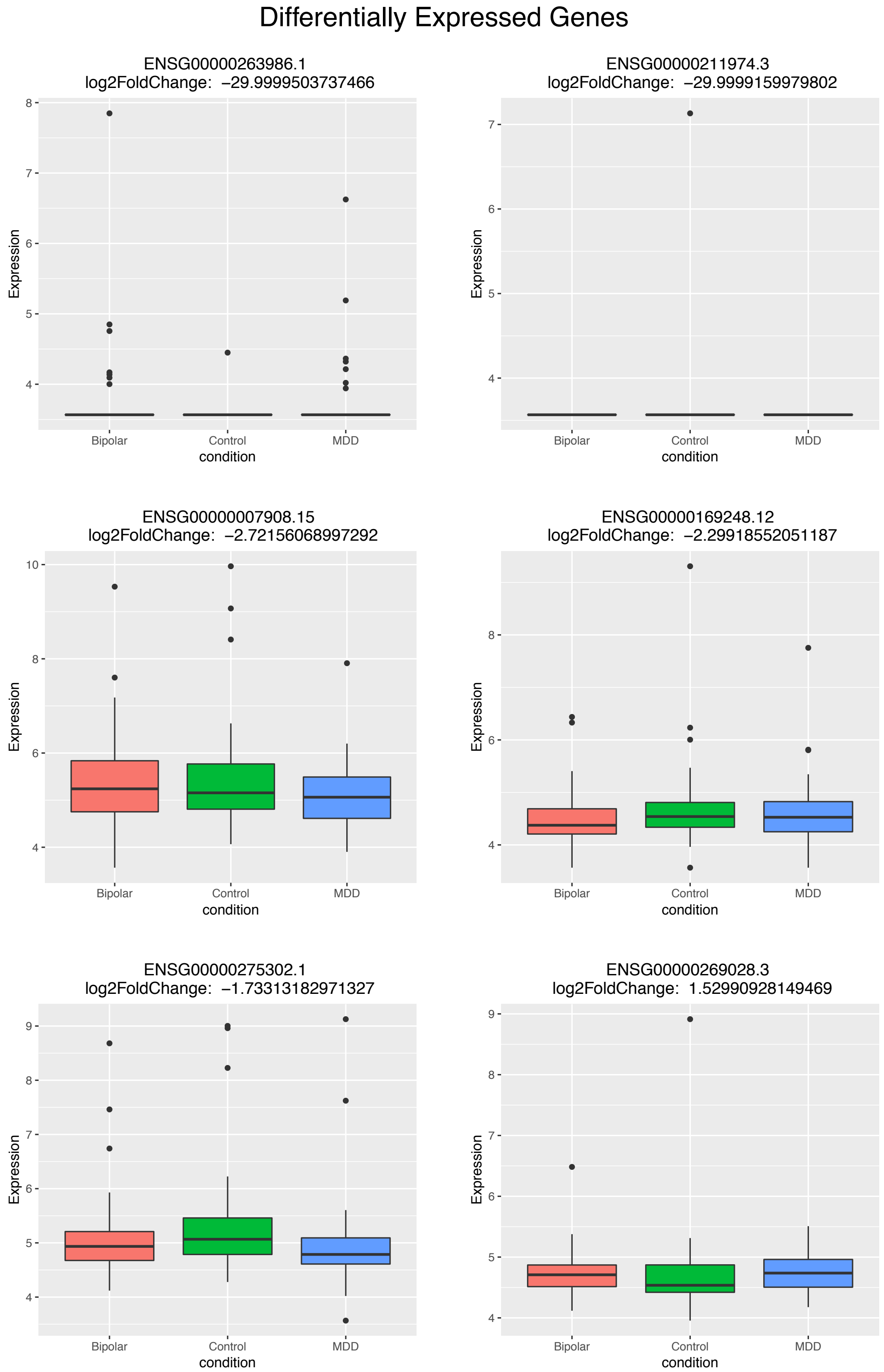


**Supplementary Figure 3**. Differential Gene Expression associated with psychiatric morbidity-&-suicide-mortality: high bipolar disorder vs low bipolar disorder & Controls. Boxplots illustrates group-specific expression mean expression values for each of the differentially expressed genes shown in Table 2B and Figure 2A. Error bars denote log-fold-change standard errors.

#### Supplementary Figure 4

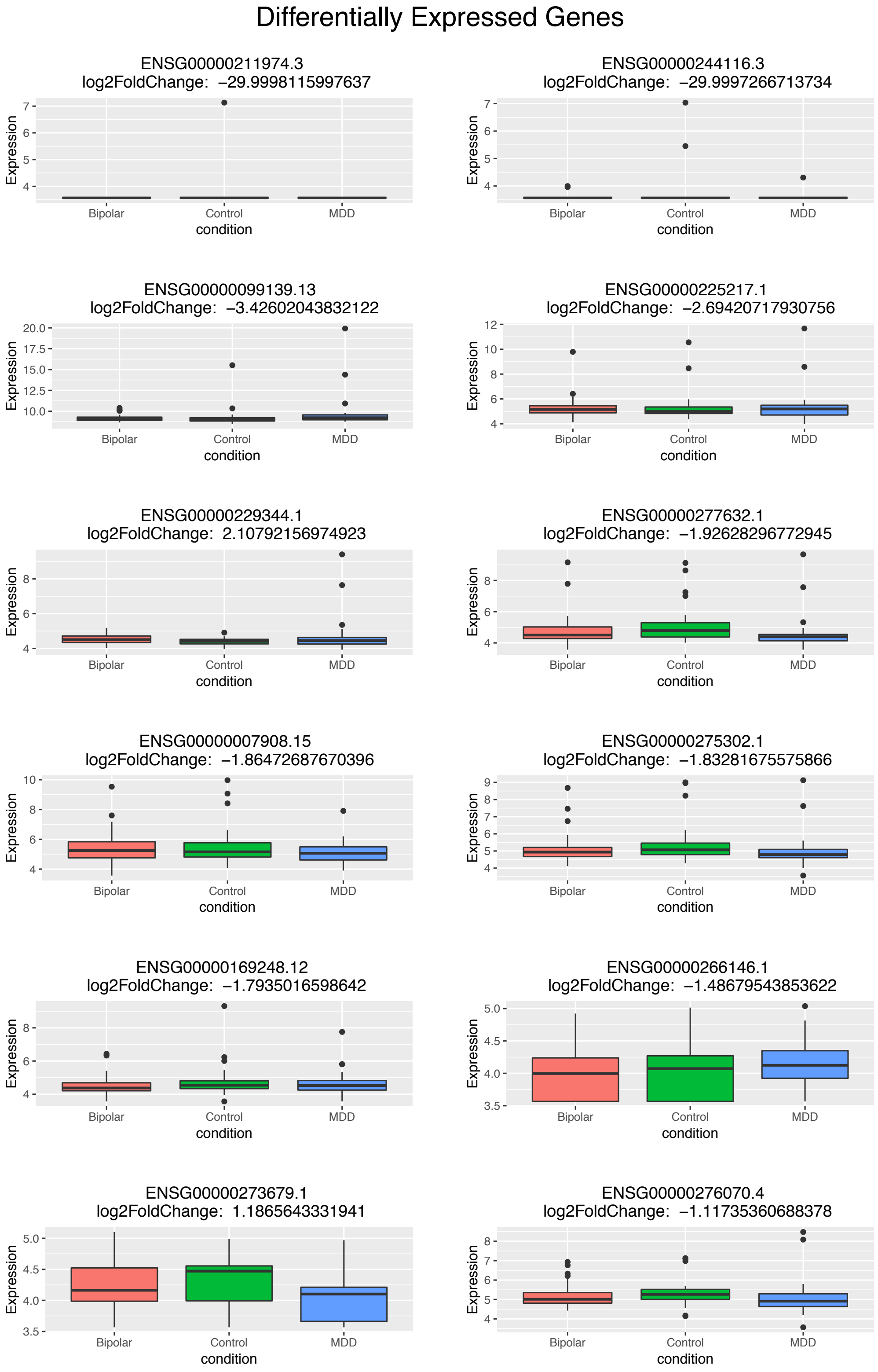


**Supplementary Figure 4**. Differential Gene Expression associated with psychiatric morbidity-&-suicide-mortality: high mood disorder combined vs low mood disorder combined vs Controls. Boxplots illustrates group-specific expression mean expression values for each of the differentially expressed genes shown in Table 2A and Figure 2A. Error bars denote log-fold-change standard errors.

#### Supplementary Figure 5

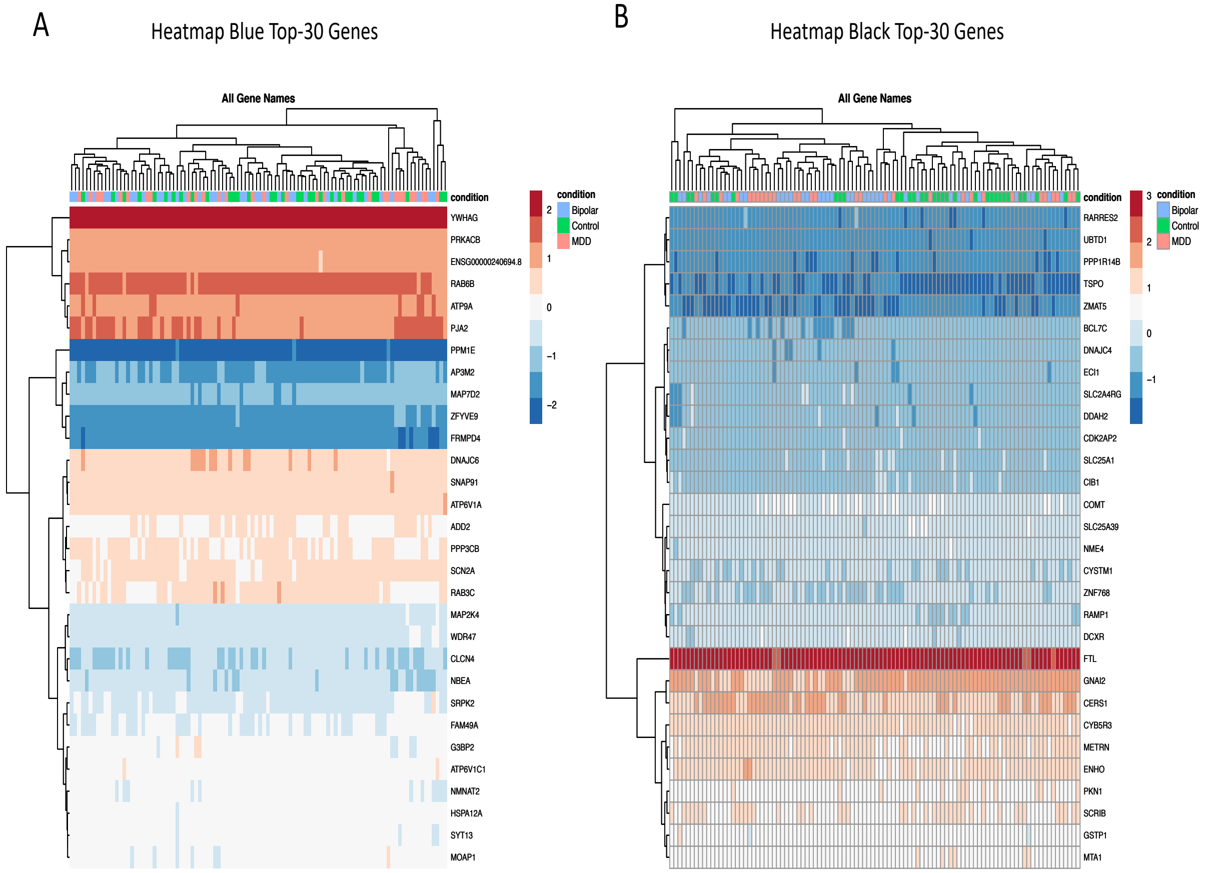


**Supplementary Figure 8**. **WGCNA Identified clinical and demographically-associated gene co-expression modules.** A). Heatmap showing the gene-expression of the 30 hub genes in the Blue module that predominantly captures genetic pathways involved in fundamental cellular signaling processes such as ATPase activity, cAMP signaling, sodium and calcium channel functions, transcriptional regulation, and neuronal development in terms of proliferation, differentiation. B) Heatmap showing the gene-expression of the 30 hub genes in the Black module that predominantly captures genetic pathways involved in innate immune functions, cellular homeostasis and metabolic regulations, dopamine DARPP32 feedback onto cAMP functions and glial cell development. Dendrograms represent sample and gene clustering by expression.

**Supplementary Table 1 (Included publications and Studies/Gray Matter Analysis Contrasts in the Imaging Meta-analysis).**

| Included | Year | 1st Author | Journal | Gray Matter Contrast Analyzed |
|---|---|---|---|---|
| False | 2005 | Adler C M | Biol Psychiatry | Bipolar > Healthy Controls |
| False | 2005 | Adler C M | Biol Psychiatry | Healthy Controls < Bipolar |
| False | 2005 | Adler C M | Biol Psychiatry | First Episode Bipolar < Multi-Episode Bipolar |
| **True** | 2007 | Chen X | The Aust. & New Zealand J. of Psych | Bipolar & Family History < Healthy Controls |
| False | 2007 | Chen X | The Aust. & New Zealand J. of Psych | Bipolar with Family History > Healthy Controls |
| **True** | 2007 | Chen X | The Aust. & New Zealand J. of Psych | Bipolar No Family History < Healthy Controls |
| **True** | 2007 | Chen X | The Aust. & New Zealand J. of Psych | Bipolar No Family History < Healthy Controls |
| False | 2007 | Chen X | The Aust. & New Zealand J. of Psych | Bipolar & Psychosis > Healthy Controls |
| **True** | 2007 | Chen X | The Aust. & New Zealand J. of Psych | Bipolar & Psychosis < Healthy Controls |
| **True** | 2007 | Chen X | The Aust. & New Zealand J. of Psych | Bipolar without Psychosis < Healthy Controls |
| **True** | 2007 | Chen X | The Aust. & New Zealand J. of Psych | Bipolar On Lithium < Healthy Controls |
| False | 2007 | Chen X | The Aust. & New Zealand J. of Psych | Bipolar Not On Lithium > Healthy Controls |
| **True** | 2007 | Chen X | The Aust. & New Zealand J. of Psych | Bipolar Not On Lithium < Healthy Controls |
| False | 2007 | Chen X | The Aust. & New Zealand J. of Psych | Bipolar & Family History > Bipolar No FH |
| False | 2007 | Chen X | The Aust. & New Zealand J. of Psych | Bipolar & Psychosis > Bipolar No Psychosis |
| False | 2007 | Chen X | The Aust. & New Zealand J. of Psych | Bipolar On Lithium > Bipolar Not On Lithium |
| False | 2007 | Chen X | The Aust. & New Zealand J. of Psych | Gray Matter-Duration of Bipolar Correlation |
| False | 2007 | Chen X | The Aust. & New Zealand J. of Psych | Gray Matter-Number of Episodes Correlation |
| **True** | 2004 | Lyoo I K | Biological Psychiatry | Healthy Controls > Bipolar Gray Matter Density |

| | | | | |
|---|---|---|---|---|
| **True** | 2009 | Almeida J R | Psychiatry Research NeuroImaging | Bipolar Disorder < Healthy Control |
| False | 2009 | Almeida J R | Psychiatry Research NeuroImaging | Male > Female Bipolar |
| **True** | 2009 | Almeida J R | Psychiatry Research NeuroImaging | Healthy Controls > Bipolar |
| False | 2009 | Almeida J R | Psychiatry Research NeuroImaging | All Males > Females |
| False | 2009 | Almeida J R | Psychiatry Research NeuroImaging | All Females > Males |
| **True** | 2009 | Bergouignan | NeuroImage | Unipolar Depressed < Healthy Controls |
| **True** | 2010 | Peng J | European Journal of Radiology | Unipolar Depressed < Healthy Controls |
| **True** | 2010 | Ha T H | Neuroscience Letters | Bipolar Disorder II < Healthy Controls |
| **True** | 2010 | Ha T H | Neuroscience Letters | Bipolar Disorder I < Healthy Controls |
| False | 2010 | Ha T H | Neuroscience Letters | Bipolar Disorder I < Bipolar Disorder II |
| False | 2010 | Ha T H | Neuroscience Letters | Onset Age-Gray Matter correlation in Bipolar II |
| **True** | 2010 | Abe O | Psychiatry Research | Unipolar Depressed < Healthy Controls |
| **True** | 2009 | Stanfield A | Bipolar Disorders | Bipolar < Healthy Controls |
| False | 2009 | Stanfield A | Bipolar Disorders | Bipolar < Healthy Controls |
| False | 2009 | Stanfield A | Bipolar Disorders | Age-GM Density correlation in Healthy Controls and Bipolar Disorder |
| **True** | 2008 | Kim M J | Psychiatry Research | Healthy Controls > Unipolar Depressed |
| **True** | 2011 | Wagner G | NeuroImage | Unipolar Depressed < Healthy Controls |
| False | 2011 | Wagner G | NeuroImage | Depressed High Risk Suicide < Healthy Controls |
| False | 2011 | Wagner G | NeuroImage | Unipolar Depressed High Risk Suicide < Unipolar Depressed Non-High Risk Suicide |
| False | 2011 | Wagner G | NeuroImage | Unipolar Depressed with Suicidal Behavior < Unipolar |

| | | | | |
|---|---|---|---|---|
| | | | | Depressed Non-High Risk Suicide |
| False | 2011 | Cui L | Neuroscience Letters | Paranoid-type Schizophrenia < Healthy Controls |
| False | 2011 | Cui L | Neuroscience Letters | Paranoid-type Schizophrenia > Healthy Controls |
| **True** | 2011 | Cui L | Neuroscience Letters | Bipolar Mania < Healthy Controls |
| False | 2011 | Cui L | Neuroscience Letters | Bipolar Mania > Healthy Controls |
| **True** | 2009 | Arnone D | European Neuropsychopharmacology | Unipolar Depression < Healthy Controls |
| **True** | 2010 | Cheng Y | Neuroscience Letters | Unipolar Depressed < Healthy Controls |
| **True** | 2010 | Cheng Y | Neuroscience Letters | Depression Score-Gray Matter Correlation |
| **True** | 2008 | Frodl T | Archives of General Psychiatry | Unipolar Depressed < Healthy Controls |
| **True** | 2010 | Hwang J | Journal of Ger. Psych and Neurology | Unipolar Depressed < Healthy Controls |
| False | 2010 | Hwang J | Journal of Ger. Psych and Neurology | Healthy Controls < Unipolar Depressed |
| False | 2010 | Hwang J | Journal of Ger. Psych and Neurology | Unipolar Depressed > Healthy Controls |
| False | 2010 | Hwang J | Journal of Ger. Psych and Neurology | Late-Onset Suicidal Depressives < Late-Onset Non-suicidal Depressives |
| False | 2010 | Hwang J | Journal of Ger. Psych and Neurology | Suicidal > Non Suicidal |
| **True** | 2010 | Li C T | NeuroImage | Non-Remitting Unipolar Depressed < Remitting Unipolar Depressives |
| False | 2010 | Li C T | NeuroImage | Remitting Unipolar Depressed > Non-remitting Unipolar Depressed |
| False | 2010 | Li C T | NeuroImage | Non-remitting Unipolar Depressed < Remitting Unipolar Depressed |
| **True** | 2011 | Salvadore G | NeuroImage | Chronic Unipolar Depressed |
| **True** | 2011 | Salvadore G | NeuroImage | Chronic Unipolar Depressed |
| **True** | 2011 | Salvadore G | NeuroImage | Remission Unipolar Depressed |

| | | | | |
|---|---|---|---|---|
| **True** | 2011 | Salvadore G | NeuroImage | Chronic Unipolar Depressed |
| **True** | 2011 | Salvadore G | NeuroImage | Chronic Unipolar Depressed |
| **True** | 2011 | Salvadore G | NeuroImage | Remission Unipolar Depressed |
| **True** | 2011 | Soriano-Mas | Biol Psychiatry | Unipolar Depressed |
| False | 2011 | Soriano-Mas | Biol Psychiatry | Healthy Controls |
| False | 2011 | Soriano-Mas | Biol Psychiatry | Unipolar Depressed-HAM-D Correlation |
| False | 2011 | Soriano-Mas | Biol Psychiatry | Unipolar Depressed-Gray Matter Correlation with Days to Remission After Treatment |
| False | 2011 | Soriano-Mas | Biol Psychiatry | Old vs. Follow-Up, Gray Matter Volume |
| False | 2011 | Soriano-Mas | Biol Psychiatry | Correlation Gray Matter Volume vs. # of Depression Episodes Between Both Scan Points |
| False | 2011 | Soriano-Mas | Biol Psychiatry | Correlation White Matter Volume vs. # of Depression Episodes Between Both Scan Points |
| False | 2011 | Soriano-Mas | Biol Psychiatry | Gray Matter Volume Decreases, Male Unipolar Depressed |
| False | 2011 | Soriano-Mas | Biol Psychiatry | White Matter Volume Decreases, Old Patients |
| False | 2010 | van Tol M J | Archives of General Psychiatry | Unipolar Depressed/CDA/ANX |
| **True** | 2010 | van Tol M J | Archives of General Psychiatry | Unipolar Depressed |
| **True** | 2010 | van Tol M J | Archives of General Psychiatry | CDA |
| False | 2010 | van Tol M J | Archives of General Psychiatry | ANX |
| False | 2010 | van Tol M J | Archives of General Psychiatry | UNIPOLAR DEPRESSED/CDA/ANX |
| **True** | 2010 | van Tol M J | Archives of General Psychiatry | Early vs. Late Onset Unipolar Depressed |
| False | 2010 | van Tol M J | Archives of General Psychiatry | ACC Reduction, Unipolar Depressed |
| False | 2010 | van Tol M J | Archives of General Psychiatry | ACC Reduction, ANX Group |

| | | | | |
|---|---|---|---|---|
| **True** | 2010 | Zou K | Biol Psychiatry | Depression |
| **True** | 2008 | Haldane M | Journal of Psychopharmacology | Bipolar |
| False | 2008 | Haldane M | Journal of Psychopharmacology | Bipolar > CS |
| False | 2008 | Haldane M | Journal of Psychopharmacology | Negative SCWT Score-Gray Matter Correlation |
| False | 2008 | Haldane M | Journal of Psychopharmacology | Negative SCWT Score-Gray Matter Correlation |
| False | 2008 | Haldane M | Journal of Psychopharmacology | Negative HSCT Score-Gray Matter Correlation |
| False | 2008 | Haldane M | Journal of Psychopharmacology | Negative HSCT Score-Gray Matter Correlation |
| **True** | 2011 | Lee H Y | Journal of Affective Disorders | Healthy Controls > Unipolar Depressed |
| False | 2011 | Lee H Y | Journal of Affective Disorders | Depression duration-Gray Matter Correlation |
| **True** | 2011 | Li M | Psychiatry Research Neuro-Imaging | Bipolar Disorder |
| **True** | 2011 | Wang F | Brain | Bipolar Disorder |
| False | 2012 | Watson D R | Behavioural Brain Research | Schizophrenia |
| False | 2012 | Watson D R | Behavioural Brain Research | Schizophrenia > sCS |
| False | 2012 | Watson D R | Behavioural Brain Research | Schizophrenia |
| False | 2012 | Watson D R | Behavioural Brain Research | Schizophrenia > sCS |
| **True** | 2012 | Watson D R | Behavioural Brain Research | Bipolar |
| False | 2012 | Watson D R | Behavioural Brain Research | Bipolar > bCS |
| False | 2012 | Watson D R | Behavioural Brain Research | Bipolar |
| **True** | 2013 | Serra-Blasco | British Journal of Psychiatry | Healthy Controls > Unipolar Depressed |
| **True** | 2013 | Serra-Blasco | British Journal of Psychiatry | Healthy Controls > Unipolar Depressed |
| **True** | 2013 | Serra-Blasco | British Journal of Psychiatry | fUnipolar Depressed > tUnipolar Depressed |
| **True** | 2012 | Shad M U | J. of Child & Adol. Psychopharm. | Healthy Controls > Unipolar Depressed |
| **True** | 2012 | Zhang X | Journal of Affective Disorders | Healthy Controls > CVD |
| **True** | 2012 | Zhang X | Journal of Affective Disorders | Healthy Controls > Unipolar Depressed |

| | | | | |
|---|---|---|---|---|
| False | 2012 | Zhang X | Journal of Affective Disorders | Unipolar Depressed > CVD |
| **True** | 2012 | Zhang X | Journal of Affective Disorders | Unipolar Depressed > Healthy Controls |
| False | 2012 | Zhang X | Journal of Affective Disorders | CES-D(Depr Scale)Score- Gray Matter Correlation |
| False | 2012 | Zhang X | Journal of Affective Disorders | Weakest Link CSQ score-Gray Matter Correlation |
| False | 2012 | Zhang X | Journal of Affective Disorders | Consequences CSQ score-Gray Matter Correlation |
| False | 2012 | Zhang X | Journal of Affective Disorders | Causal Attributions CSQ-Gray Matter Correlation |
| False | 2012 | Zhang X | Journal of Affective Disorders | Gray Matter Volume, Main Effect of Group |
| **True** | 2013 | Arnone D | Molecular Psychiatry | Unipolar Depressed < Healthy controls |
| False | 2013 | Arnone D | Molecular Psychiatry | Unipolar Depressed < Remitted Unipolar Depressed |
| False | 2013 | Arnone D | Molecular Psychiatry | Currently Unipolar Depressed > Healthy controls |
| False | 2013 | Arnone D | Molecular Psychiatry | Remitted Unipolar Depressed > Healthy Controls |
| False | 2013 | Arnone D | Molecular Psychiatry | Effect of medication in remitted patients |
| False | 2013 | Arnone D | Molecular Psychiatry | cUinpolar symptoms-Grey matter Correlation |
| False | 2013 | Grieve S M | NeuroImage | Unipolar Depressed > Controls, Cortical thickness |
| **True** | 2013 | Grieve S M | NeuroImage | Healthy Controls > Unipolar Depressed |
| **True** | 2013 | Grieve S M | NeuroImage | Healthy Controls > Unipolar Depressed |
| **True** | 2014 | Stratmann | PLoS One | Healthy Controls > Unipolar Depressed |
| **True** | 2014 | Stratmann | PLoS One | Healthy Controls > Current Unipolar Depressed |
| False | 2014 | Stratmann M | PLoS One | Depressed Episodes-Grey Matter Correlation |
| **True** | 2014 | Guo W | Prog. In Neuro-Psychopharm. & Biological Psychiatry | Unipolar Depressed < Healthy Controls |

| | | | | |
|---|---|---|---|---|
| **True** | 2014 | Guo W | Prog. In Neuro-Psychopharm. & Biological Psychiatry | 1st episode Unipolar Depressed < Healthy Controls |
| **True** | 2014 | Guo W | Prog. In Neuro-Psychopharm. & Biological Psychiatry | Recurrent Unipolar Depressed < Healthy Controls |
| **True** | 2014 | Tang L R | Psychiatry Research NeuroImaging | Healthy Controls > Bipolar I |
| False | 2014 | Tang L R | Psychiatry Research NeuroImaging | Healthy Controls < Bipolar I |
| False | 2014 | Tang L R | Psychiatry Research NeuroImaging | Brain regions with statistically significant volume differences, both groups |
| **True** | 2010 | Tost H | Journal of Affective Disorders | Healthy Controls > Bipolar |
| **True** | 2010 | Tost H | Journal of Affective Disorders | Healthy Controls > Bipolar |
| False | 2010 | Tost H | Journal of Affective Disorders | Healthy Controls > Bipolar |
| False | 2010 | Tost H | Journal of Affective Disorders | Bipolar > Healthy Controls |
| False | 2010 | Tost H | Journal of Affective Disorders | Bipolar > Healthy Controls |
| **True** | 2011 | Gong Q | NeuroImage | Healthy Controls > Refractory Depression |
| False | 2011 | Gong Q | NeuroImage | Refractory Depression > Healthy Controls |
| **True** | 2011 | Gong Q | NeuroImage | Healthy Controls > Non-Refractory Depression |
| False | 2011 | Gong Q | NeuroImage | NDD > Healthy Controls |
| False | 2011 | Gong Q | NeuroImage | Healthy Controls > RDD |
| False | 2011 | Gong Q | NeuroImage | RDD > Healthy Controls |
| False | 2011 | Gong Q | NeuroImage | Healthy Controls > NDD |
| False | 2011 | Gong Q | NeuroImage | NDD > Healthy Controls |
| False | 2015 | Cai Y | Neuroscience Bulletin | Main effect of group, gray matter |
| **True** | 2015 | Cai Y | Neuroscience Bulletin | Bipolar I < Healthy Controls |
| **True** | 2015 | Cai Y | Neuroscience Bulletin | Unipolar Depressed < Healthy Controls |
| False | 2015 | Cai Y | Neuroscience Bulletin | Bipolar I < Unipolar Depressed |
| **True** | 2013 | Kim D | Journal of Affective Disorders | Bipolar < Healthy Controls |

| | | | | |
|---|---|---|---|---|
| False | 2015 | Saricicek A | Journal of Affective Disorders | Group Gray matter differences |
| False | 2015 | Saricicek A | Journal of Affective Disorders | Bipolar I > Healthy Controls |
| **True** | 2015 | Saricicek A | Journal of Affective Disorders | Bipolar I < Healthy Controls |
| False | 2015 | Saricicek A | Journal of Affective Disorders | Healthy first-degree relatives > Healthy Controls |
| False | 2015 | Saricicek A | Journal of Affective Disorders | Healthy first-degree relatives < Healthy Controls |
| False | 2014 | Redlich R | Archives of General Psychiatry | Munster Sample |
| False | 2014 | Redlich R | Archives of General Psychiatry | Pittsburgh Sample |
| False | 2014 | Redlich R | Archives of General Psychiatry | Combined Sample Pittsburgh & Munster Samples |
| **True** | 2014 | Redlich R | Archives of General Psychiatry | Unipolar Depressed < Healthy Controls |
| **True** | 2014 | Redlich R | Archives of General Psychiatry | Bipolar < Healthy Controls |
| False | 2014 | Redlich R | Archives of General Psychiatry | Bipolar < Unipolar Depressed Munster Sample |
| False | 2014 | Redlich R | Archives of General Psychiatry | Unipolar Depressed < Bipolar Munster Sample |
| False | 2014 | Redlich R | Archives of General Psychiatry | Bipolar < Unipolar Depressed Pittsburgh Sample |
| False | 2014 | Redlich R | Archives of General Psychiatry | Unipolar Depressed < Bipolar Pittsburgh Sample |
| False | 2014 | Redlich R | Archives of General Psychiatry | Bipolar < Unipolar Depressed Combined Sample |
| False | 2014 | Redlich R | Archives of General Psychiatry | Unipolar Depressed < Bipolar Combined Sample |
| False | 2014 | Redlich R | Archives of General Psychiatry | Effect of medication load + clinical course of illness on gray matter, MD patients vs Bipolar |
| False | 2014 | Redlich R | Archives of General Psychiatry | Gray matter volume-Unipolar Depressed illness duration Correlation |

| | | | | |
|---|---|---|---|---|
| False | 2014 | Redlich R | Archives of General Psychiatry | Gray matter volume-Unipolar Depressed illness duration, Munster sample |
| False | 2014 | Redlich R | Archives of General Psychiatry | Gray matter volume-Unipolar Depressed illness duration, Pittsburgh sample |
| False | 2012 | Adleman N | Journal of Child Psychology and Psychiatry | Bipolar > Healthy Controls and SMD |
| False | 2012 | Adleman N | Journal of Child Psychology and Psychiatry | Bipolar Disorder > SMD |
| **True** | 2012 | Adleman N | Journal of Child Psychology and Psychiatry | Healthy Controls > Bipolar Disorder |
| False | 2012 | Adleman N | Journal of Child Psychology and Psychiatry | 2 year scan > Initial scan; Bipolar group |
| **True** | 2016 | Alonso-Lana | PLoS ONE | Cognitively preserved patients < Healthy Controls |
| False | 2016 | Alonso-Lana | PLoS ONE | Cognitively preserved patients < Healthy Controls |
| False | 2015 | Lai C H | Journal of Affective Disorders | Healthy Controls > Panic Disorder |
| **True** | 2015 | Lai C H | Journal of Affective Disorders | Healthy Controls > Unipolar Depressed |
| False | 2015 | Lai C H | Journal of Affective Disorders | Panic Disorder > Unipolar Depressed |
| **True** | 2012 | Singh M K | Bipolar Disorders | Bipolar I < Healthy Controls |
| **True** | 2012 | Singh M K | Bipolar Disorders | Bipolar < Healthy Controls |

**Supplementary Table 2**

| **Controls > MDD Simple Comparison** |
|---|

| **Ensemble ID** | **baseMean** | **log2FoldChange** | **lfcSE** | **stat** | **pvalue** | **padj** |
|---|---|---|---|---|---|---|
| ENSG00000121207.11 | 197.2273518 | 0.912906057 | 0.16032 | 5.694347951 | 1.24E-08 | 0.00012742 |
| ENSG00000160606.10 | 69.57779939 | 1.035594387 | 0.19829 | 5.22252558 | 1.76E-07 | 0.000454 |
| ENSG00000175287.18 | 256.20944 | 0.864415678 | 0.16424 | 5.26297854 | 1.42E-07 | 0.000454 |
| ENSG00000176244.6 | 779.0974012 | 0.812075787 | 0.15888 | 5.111300328 | 3.20E-07 | 0.00054866 |
| ENSG00000214456.8 | 187.0730578 | 0.794324426 | 0.15524 | 5.116793583 | 3.11E-07 | 0.00054866 |
| ENSG00000124440.15 | 606.769243 | 0.941920116 | 0.19051 | 4.944165422 | 7.65E-07 | 0.00102124 |
| ENSG00000147119.3 | 251.7565669 | 0.931770988 | 0.19099 | 4.878552296 | 1.07E-06 | 0.00102124 |
| ENSG00000166292.11 | 99.84988779 | 0.889665426 | 0.1808 | 4.920679692 | 8.62E-07 | 0.00102124 |
| ENSG00000279103.1 | 99.94132755 | 0.912942009 | 0.18653 | 4.894263276 | 9.87E-07 | 0.00102124 |
| ENSG00000056487.15 | 125.6707575 | 0.757104683 | 0.16074 | 4.710021333 | 2.48E-06 | 0.00106187 |
| ENSG00000111087.9 | 36.37791249 | 1.082827773 | 0.22782 | 4.752943591 | 2.00E-06 | 0.00106187 |
| ENSG00000132164.9 | 965.8191898 | 0.639572988 | 0.13417 | 4.76672186 | 1.87E-06 | 0.00106187 |
| ENSG00000138759.17 | 756.8359079 | -0.621747945 | 0.12873 | -4.829887631 | 1.37E-06 | 0.00106187 |
| ENSG00000165478.6 | 3937.245506 | 0.99043693 | 0.20437 | 4.846283464 | 1.26E-06 | 0.00106187 |
| ENSG00000168209.4 | 815.1293443 | 0.992762628 | 0.20975 | 4.733050352 | 2.21E-06 | 0.00106187 |
| ENSG00000177133.10 | 738.3511464 | 0.907612217 | 0.19099 | 4.752027849 | 2.01E-06 | 0.00106187 |
| ENSG00000246022.2 | 61.45792813 | 0.761495297 | 0.15919 | 4.783495044 | 1.72E-06 | 0.00106187 |
| ENSG00000255690.2 | 1413.820761 | 0.852295666 | 0.17993 | 4.736860733 | 2.17E-06 | 0.00106187 |
| ENSG00000280087.1 | 181.6095253 | 0.784637368 | 0.16505 | 4.753926066 | 2.00E-06 | 0.00106187 |
| ENSG00000111405.8 | 50.70506442 | 0.770791032 | 0.1672 | 4.6100377 | 4.03E-06 | 0.00109008 |
| ENSG00000134508.12 | 1958.795414 | 0.692949658 | 0.15027 | 4.611268395 | 4.00E-06 | 0.00109008 |
| ENSG00000137434.11 | 13.49298633 | 0.734441682 | 0.15926 | 4.611532137 | 4.00E-06 | 0.00109008 |
| ENSG00000211592.8 | 16.70325767 | -2.66188589 | 0.57006 | -4.669489567 | 3.02E-06 | 0.00109008 |
| ENSG00000239282.7 | 138.4067896 | 0.740203148 | 0.16037 | 4.615705028 | 3.92E-06 | 0.00109008 |
| ENSG00000259823.5 | 182.3106437 | -0.669209741 | 0.14509 | -4.612286033 | 3.98E-06 | 0.00109008 |
| ENSG00000278709.1 | 21.84241556 | -0.666984434 | 0.14399 | -4.632104909 | 3.62E-06 | 0.00109008 |
| ENSG00000159423.16 | 1426.850524 | 0.82323545 | 0.17927 | 4.592096602 | 4.39E-06 | 0.00114466 |
| ENSG00000129007.14 | 36.5005764 | 1.018111508 | 0.22473 | 4.530279197 | 5.89E-06 | 0.00130192 |
| ENSG00000154930.14 | 1415.939276 | 0.811596969 | 0.17903 | 4.533290306 | 5.81E-06 | 0.00130192 |
| ENSG00000165795.22 | 19499.20092 | 0.679472214 | 0.15005 | 4.528259505 | 5.95E-06 | 0.00130192 |
| ENSG00000167676.4 | 166.0240233 | 1.114639029 | 0.24587 | 4.533481331 | 5.80E-06 | 0.00130192 |
| ENSG00000183773.15 | 1468.88116 | 0.607680351 | 0.13405 | 4.533378318 | 5.80E-06 | 0.00130192 |
| ENSG00000271447.5 | 315.5409592 | 0.80838449 | 0.17833 | 4.533126353 | 5.81E-06 | 0.00130192 |

| | | | | | | |
|---|---|---|---|---|---|---|
| ENSG00000121570.12 | 236.5686399 | 1.378421234 | 0.30481 | 4.522206959 | 6.12E-06 | 0.00131181 |
| ENSG00000166033.11 | 4464.65292 | 0.621604376 | 0.1378 | 4.510914758 | 6.45E-06 | 0.00132828 |
| ENSG00000088826.17 | 724.5980016 | 0.659384469 | 0.14741 | 4.47320281 | 7.71E-06 | 0.0014804 |
| ENSG00000154856.12 | 1024.659047 | 0.673911661 | 0.15071 | 4.47143604 | 7.77E-06 | 0.0014804 |
| ENSG00000167315.17 | 669.8439443 | 0.62334091 | 0.13927 | 4.475921583 | 7.61E-06 | 0.0014804 |
| ENSG00000139800.8 | 131.7381248 | 0.628041896 | 0.14064 | 4.465569984 | 7.99E-06 | 0.00149024 |
| ENSG00000272189.1 | 176.4028167 | 0.91382829 | 0.20497 | 4.458441216 | 8.26E-06 | 0.00149024 |
| ENSG00000073756.11 | 512.9418063 | -1.198423659 | 0.26918 | -4.452155349 | 8.50E-06 | 0.00149622 |
| ENSG00000095917.13 | 12.2472475 | 1.615858283 | 0.36357 | 4.444459008 | 8.81E-06 | 0.00149622 |
| ENSG00000132692.18 | 5194.371495 | 0.626566604 | 0.14117 | 4.438328478 | 9.07E-06 | 0.00149622 |
| ENSG00000157833.12 | 852.7857104 | 0.778270948 | 0.17534 | 4.438631745 | 9.05E-06 | 0.00149622 |
| ENSG00000214353.7 | 138.2393918 | 0.737203309 | 0.16657 | 4.42579536 | 9.61E-06 | 0.0015254 |
| ENSG00000124171.8 | 282.9921114 | 0.602099597 | 0.13686 | 4.399465257 | 1.09E-05 | 0.00164197 |
| ENSG00000263146.2 | 320.5807765 | 0.593334648 | 0.13483 | 4.400761292 | 1.08E-05 | 0.00164197 |
| ENSG00000100033.16 | 1691.486571 | 0.787924933 | 0.1804 | 4.367640699 | 1.26E-05 | 0.00169577 |
| ENSG00000103740.9 | 1488.83305 | 0.861252384 | 0.19741 | 4.362734213 | 1.28E-05 | 0.00169577 |
| ENSG00000128311.13 | 189.6089781 | 0.629569794 | 0.14366 | 4.38235733 | 1.17E-05 | 0.00169577 |
| ENSG00000196071.4 | 86.23646088 | -0.703781049 | 0.16071 | -4.379174441 | 1.19E-05 | 0.00169577 |
| ENSG00000275294.4 | 11.26306985 | 0.767843225 | 0.176 | 4.362813955 | 1.28E-05 | 0.00169577 |
| ENSG00000279175.1 | 174.6828851 | 0.952260472 | 0.21842 | 4.359764298 | 1.30E-05 | 0.00169577 |
| ENSG00000279474.1 | 28.55098334 | 0.794937356 | 0.18232 | 4.360108364 | 1.30E-05 | 0.00169577 |
| ENSG00000260084.1 | 26.09702632 | 0.610429272 | 0.14027 | 4.351735462 | 1.35E-05 | 0.00169694 |
| ENSG00000160678.11 | 1588.497859 | 0.698079895 | 0.16059 | 4.346836742 | 1.38E-05 | 0.00171213 |
| ENSG00000138696.10 | 709.7537742 | 0.851444421 | 0.19621 | 4.339370863 | 1.43E-05 | 0.00172966 |
| ENSG00000225194.2 | 126.8388738 | 0.650156918 | 0.15037 | 4.32358944 | 1.54E-05 | 0.0018366 |
| ENSG00000251372.5 | 64.1816606 | 0.726279915 | 0.16833 | 4.314662965 | 1.60E-05 | 0.00184208 |
| ENSG00000130876.11 | 242.5715949 | 0.908811667 | 0.21203 | 4.286283775 | 1.82E-05 | 0.0019887 |
| ENSG00000164089.8 | 2420.394698 | 0.9752965 | 0.228 | 4.27766074 | 1.89E-05 | 0.00204554 |
| ENSG00000163395.16 | 171.5377101 | 1.01124833 | 0.23675 | 4.271398808 | 1.94E-05 | 0.00206046 |
| ENSG00000068078.17 | 3998.838972 | 0.999833773 | 0.23422 | 4.268767442 | 1.97E-05 | 0.00206364 |
| ENSG00000016391.10 | 628.9316715 | 0.678425059 | 0.15939 | 4.256263544 | 2.08E-05 | 0.00212744 |
| ENSG00000099139.13 | 842.875837 | 1.12900565 | 0.26581 | 4.2474048 | 2.16E-05 | 0.00213953 |
| ENSG00000164292.12 | 2192.047637 | 0.668110501 | 0.15729 | 4.247628387 | 2.16E-05 | 0.00213953 |
| ENSG00000100427.15 | 4774.815906 | 0.837386796 | 0.19749 | 4.24013481 | 2.23E-05 | 0.00214404 |
| ENSG00000143416.20 | 658.1302846 | 0.723181446 | 0.17053 | 4.240722092 | 2.23E-05 | 0.00214404 |
| ENSG00000178814.16 | 198.8433615 | 0.646229316 | 0.15247 | 4.238466008 | 2.25E-05 | 0.00214404 |
| ENSG00000119927.13 | 802.1382448 | 0.807335851 | 0.19094 | 4.228131663 | 2.36E-05 | 0.00221538 |
| ENSG00000125285.5 | 345.832117 | 0.774295145 | 0.18318 | 4.226982138 | 2.37E-05 | 0.00221538 |
| ENSG00000092621.11 | 1475.086995 | 0.658608218 | 0.1561 | 4.21912422 | 2.45E-05 | 0.00223311 |

| | | | | | | |
|---|---|---|---|---|---|---|
| ENSG00000115507.9 | 71.06443723 | 0.738682353 | 0.17505 | 4.219791337 | 2.45E-05 | 0.00223311 |
| ENSG00000181449.3 | 1684.839673 | 0.77430641 | 0.18388 | 4.210988472 | 2.54E-05 | 0.00227481 |
| ENSG00000253661.1 | 54.41791222 | 0.658541428 | 0.15632 | 4.212740374 | 2.52E-05 | 0.00227481 |
| ENSG00000167601.11 | 1081.028645 | 0.816203791 | 0.19402 | 4.206767038 | 2.59E-05 | 0.00229548 |
| ENSG00000182103.4 | 580.9952976 | 0.69016823 | 0.16413 | 4.205047206 | 2.61E-05 | 0.00229548 |
| ENSG00000101144.12 | 795.3251435 | 0.668741335 | 0.15962 | 4.18970749 | 2.79E-05 | 0.00233928 |
| ENSG00000266964.5 | 1091.595355 | 0.602743999 | 0.14409 | 4.18312241 | 2.88E-05 | 0.00233928 |
| ENSG00000002933.7 | 211.0700635 | 0.743675905 | 0.17839 | 4.168733595 | 3.06E-05 | 0.00234438 |
| ENSG00000119711.12 | 3819.45159 | 0.604914193 | 0.1452 | 4.166081772 | 3.10E-05 | 0.00234438 |
| ENSG00000154319.14 | 798.0289891 | 0.701869151 | 0.16839 | 4.168166012 | 3.07E-05 | 0.00234438 |
| ENSG00000203414.2 | 98.92332838 | 1.108763349 | 0.26581 | 4.171211891 | 3.03E-05 | 0.00234438 |
| ENSG00000255503.1 | 16.05253308 | 0.648265311 | 0.15556 | 4.167247844 | 3.08E-05 | 0.00234438 |
| ENSG00000171517.5 | 63.01099453 | 0.931041855 | 0.2243 | 4.150884765 | 3.31E-05 | 0.00245154 |
| ENSG00000106003.12 | 245.7342754 | 0.744894137 | 0.18104 | 4.1144664 | 3.88E-05 | 0.00273487 |
| ENSG00000176399.3 | 13.16853562 | 0.83833701 | 0.20391 | 4.111357156 | 3.93E-05 | 0.00275311 |
| ENSG00000182902.13 | 1982.59053 | 0.660328265 | 0.16085 | 4.105271493 | 4.04E-05 | 0.00277008 |
| ENSG00000183580.9 | 396.0072939 | 0.6059737 | 0.14809 | 4.091819775 | 4.28E-05 | 0.00285955 |
| ENSG00000121742.16 | 897.2620946 | 1.011184292 | 0.24725 | 4.089653714 | 4.32E-05 | 0.00286776 |
| ENSG00000130203.9 | 10421.97846 | 0.888667423 | 0.21752 | 4.085393964 | 4.40E-05 | 0.00286889 |
| ENSG00000185960.13 | 63.91012304 | 1.39584739 | 0.34169 | 4.085113683 | 4.41E-05 | 0.00286889 |
| ENSG00000185960.13_PAR_Y | 63.91012304 | 1.39584739 | 0.34169 | 4.085113683 | 4.41E-05 | 0.00286889 |
| ENSG00000226191.3 | 15.85430166 | -0.883535025 | 0.2165 | -4.081062567 | 4.48E-05 | 0.00288287 |
| ENSG00000228288.6 | 32.9407344 | 0.607452047 | 0.14895 | 4.078274275 | 4.54E-05 | 0.00289953 |
| ENSG00000043355.11 | 247.2893564 | 0.654853198 | 0.16119 | 4.062596895 | 4.85E-05 | 0.0029208 |
| ENSG00000105854.12 | 2458.660374 | 0.629183686 | 0.15482 | 4.063905003 | 4.83E-05 | 0.0029208 |
| ENSG00000112333.11 | 501.4092867 | 0.669098963 | 0.16465 | 4.063868172 | 4.83E-05 | 0.0029208 |
| ENSG00000126778.8 | 15.72250869 | 0.914637626 | 0.22474 | 4.06984266 | 4.70E-05 | 0.0029208 |
| ENSG00000179761.11 | 178.3330126 | 0.691766184 | 0.17006 | 4.067884385 | 4.74E-05 | 0.0029208 |
| ENSG00000254245.2 | 232.4474179 | 0.879774821 | 0.21627 | 4.067876075 | 4.74E-05 | 0.0029208 |
| ENSG00000259065.1 | 15.44445792 | 0.856526678 | 0.21059 | 4.067299019 | 4.76E-05 | 0.0029208 |
| ENSG00000281453.1 | 120.2169086 | 0.597651977 | 0.14742 | 4.053948912 | 5.04E-05 | 0.00295251 |
| ENSG00000018625.14 | 18477.44067 | 0.745858629 | 0.18427 | 4.047674659 | 5.17E-05 | 0.00299011 |
| ENSG00000111907.20 | 949.1766876 | 0.632779447 | 0.15647 | 4.044089898 | 5.25E-05 | 0.00300023 |
| ENSG00000161509.13 | 1150.281482 | 0.841509024 | 0.20814 | 4.042967603 | 5.28E-05 | 0.00300023 |
| ENSG00000197921.5 | 146.6547255 | 1.016729392 | 0.25142 | 4.043948737 | 5.26E-05 | 0.00300023 |
| ENSG00000198695.2 | 29121.02745 | 0.810562992 | 0.20065 | 4.039626539 | 5.35E-05 | 0.00302657 |
| ENSG00000101198.14 | 568.7849498 | 0.698285287 | 0.17307 | 4.034664297 | 5.47E-05 | 0.00303361 |
| ENSG00000135094.10 | 151.5535099 | 0.854189719 | 0.21198 | 4.029500923 | 5.59E-05 | 0.00304652 |
| ENSG00000164050.12 | 2508.084726 | 0.657880416 | 0.16328 | 4.029221006 | 5.60E-05 | 0.00304652 |

| | | | | | | |
|---|---|---|---|---|---|---|
| ENSG00000101439.8 | 17007.29606 | 0.687486857 | 0.17131 | 4.013199498 | 5.99E-05 | 0.00321002 |
| ENSG00000187091.13 | 327.7333023 | 0.667972733 | 0.16653 | 4.01103285 | 6.05E-05 | 0.00322284 |
| ENSG00000256463.8 | 322.554474 | 0.722992939 | 0.18051 | 4.005284447 | 6.19E-05 | 0.00328521 |
| ENSG00000150893.10 | 146.987741 | 0.831490123 | 0.20775 | 4.002402481 | 6.27E-05 | 0.00330043 |
| ENSG00000152990.13 | 1010.172519 | 0.693787756 | 0.17342 | 4.000641971 | 6.32E-05 | 0.00330043 |
| ENSG00000100979.14 | 1009.93884 | 0.657428374 | 0.16622 | 3.955191147 | 7.65E-05 | 0.00381959 |
| ENSG00000130055.13 | 135.9078751 | 0.633690308 | 0.16075 | 3.941976248 | 8.08E-05 | 0.00384947 |
| ENSG00000134569.9 | 2250.171729 | 0.713490028 | 0.18056 | 3.951458695 | 7.77E-05 | 0.00384947 |
| ENSG00000170075.8 | 2953.556585 | 0.841145294 | 0.21316 | 3.946048773 | 7.95E-05 | 0.00384947 |
| ENSG00000170370.11 | 723.897125 | 0.65095723 | 0.16503 | 3.944390722 | 8.00E-05 | 0.00384947 |
| ENSG00000107317.11 | 11635.75915 | 0.615252526 | 0.15621 | 3.938689607 | 8.19E-05 | 0.00388459 |
| ENSG00000279672.1 | 174.9131622 | 0.706079058 | 0.17949 | 3.933829765 | 8.36E-05 | 0.00392936 |
| ENSG00000181856.14 | 62.75413093 | 0.828849976 | 0.21113 | 3.925786183 | 8.64E-05 | 0.00398972 |
| ENSG00000248713.1 | 28.08973367 | -0.89323771 | 0.22753 | -3.925717129 | 8.65E-05 | 0.00398972 |
| ENSG00000165474.5 | 32.47352177 | 1.132234445 | 0.28869 | 3.922036336 | 8.78E-05 | 0.00401517 |
| ENSG00000146250.6 | 180.7431686 | 0.936059821 | 0.23926 | 3.912307402 | 9.14E-05 | 0.00414236 |
| ENSG00000089335.20 | 3003.435856 | 0.678626042 | 0.17414 | 3.897095923 | 9.74E-05 | 0.00435971 |
| ENSG00000236255.1 | 1130.596419 | 1.605895886 | 0.41264 | 3.891743426 | 9.95E-05 | 0.00443302 |
| ENSG00000179399.14 | 368.3618622 | 0.653915423 | 0.16824 | 3.886796875 | 0.000102 | 0.00449886 |
| ENSG00000267385.1 | 23.34444481 | 1.027787327 | 0.26448 | 3.886072141 | 0.000102 | 0.00449886 |
| ENSG00000144908.13 | 1325.601343 | 0.821804597 | 0.21163 | 3.883248795 | 0.000103 | 0.004532 |
| ENSG00000153446.15 | 231.0960163 | 0.743443572 | 0.19159 | 3.880470902 | 0.000104 | 0.00456457 |
| ENSG00000227640.2 | 164.5085386 | 0.656893763 | 0.16935 | 3.878838252 | 0.000105 | 0.00457584 |
| ENSG00000177045.7 | 70.26144593 | 0.810146887 | 0.20919 | 3.872819752 | 0.000108 | 0.00467057 |
| ENSG00000128602.9 | 181.3845846 | 0.718590834 | 0.18593 | 3.864831267 | 0.000111 | 0.00478572 |
| ENSG00000064655.18 | 152.3828714 | 0.766589998 | 0.19877 | 3.856763066 | 0.000115 | 0.00492579 |
| ENSG00000169006.6 | 578.5475956 | 0.841871133 | 0.21841 | 3.854509092 | 0.000116 | 0.00493183 |
| ENSG00000114315.3 | 358.0531148 | 0.882259958 | 0.22945 | 3.845183382 | 0.00012 | 0.00505223 |
| ENSG00000105852.10 | 30.10673412 | 0.749960838 | 0.19566 | 3.832917221 | 0.000127 | 0.00514417 |
| ENSG00000100767.15 | 535.6678649 | 0.775702264 | 0.20258 | 3.829176339 | 0.000129 | 0.00515167 |
| ENSG00000274956.2 | 1748.869071 | 0.650802388 | 0.16997 | 3.828972904 | 0.000129 | 0.00515167 |
| ENSG00000115380.19 | 886.5194508 | 0.921200039 | 0.24066 | 3.827819295 | 0.000129 | 0.00515581 |
| ENSG00000204347.3 | 59.81884562 | 0.787335669 | 0.20578 | 3.826113293 | 0.00013 | 0.00517161 |
| ENSG00000261026.1 | 23.76470883 | -0.679090259 | 0.17781 | -3.819275303 | 0.000134 | 0.0052562 |
| ENSG00000134595.8 | 21.38719696 | 0.716620057 | 0.18833 | 3.805186645 | 0.000142 | 0.00539972 |
| ENSG00000125144.13 | 493.4401699 | 0.751133524 | 0.19759 | 3.801532653 | 0.000144 | 0.0054397 |
| ENSG00000149090.11 | 680.6047309 | 0.64464161 | 0.16954 | 3.802213376 | 0.000143 | 0.0054397 |
| ENSG00000129244.8 | 9124.929529 | 0.696102341 | 0.18334 | 3.796689353 | 0.000147 | 0.00550654 |
| ENSG00000137285.9 | 5706.905134 | 0.737859184 | 0.19453 | 3.793108377 | 0.000149 | 0.00556628 |

| | | | | | | |
|---|---|---|---|---|---|---|
| ENSG00000140022.9 | 789.8380315 | 0.729272727 | 0.1926 | 3.786505656 | 0.000153 | 0.00564083 |
| ENSG00000146648.16 | 723.0515938 | 0.661790817 | 0.17479 | 3.786216227 | 0.000153 | 0.00564083 |
| ENSG00000089472.16 | 496.1220748 | 0.690815589 | 0.1828 | 3.779166829 | 0.000157 | 0.00574106 |
| ENSG00000109062.10 | 1045.580721 | 0.701924738 | 0.18595 | 3.774793337 | 0.00016 | 0.00574106 |
| ENSG00000135821.17 | 35484.82158 | 0.737413768 | 0.19535 | 3.774810736 | 0.00016 | 0.00574106 |
| ENSG00000233850.1 | 12.06497577 | 0.696103577 | 0.18431 | 3.776810006 | 0.000159 | 0.00574106 |
| ENSG00000272338.2 | 39.11132539 | 0.585317571 | 0.15515 | 3.77251609 | 0.000162 | 0.00574789 |
| ENSG00000273091.1 | 19.30018051 | 0.651998032 | 0.1729 | 3.770862787 | 0.000163 | 0.00574789 |
| ENSG00000081248.10 | 51.15721782 | -0.601551476 | 0.16001 | -3.759351775 | 0.00017 | 0.00594695 |
| ENSG00000170425.3 | 160.9038544 | 0.650389852 | 0.17356 | 3.747285782 | 0.000179 | 0.00615379 |
| ENSG00000068976.13 | 544.8238665 | 0.716892538 | 0.19157 | 3.742118519 | 0.000182 | 0.00617595 |
| ENSG00000127249.14 | 568.9629715 | 0.739629186 | 0.19756 | 3.743758747 | 0.000181 | 0.00617595 |
| ENSG00000163346.16 | 2517.992154 | 0.722684516 | 0.19296 | 3.745291022 | 0.00018 | 0.00617595 |
| ENSG00000185432.11 | 2388.742888 | 0.672365031 | 0.1799 | 3.737383404 | 0.000186 | 0.00625071 |
| ENSG00000136235.15 | 568.0418348 | 0.730863019 | 0.19587 | 3.731378941 | 0.00019 | 0.00631762 |
| ENSG00000170989.8 | 1279.36382 | 0.770215123 | 0.20639 | 3.731833547 | 0.00019 | 0.00631762 |
| ENSG00000267014.5 | 34.81448206 | 0.676846983 | 0.18133 | 3.732637698 | 0.000189 | 0.00631762 |
| ENSG00000169439.11 | 1026.253355 | 0.668147382 | 0.17916 | 3.729239012 | 0.000192 | 0.00633364 |
| ENSG00000238230.1 | 20.42441878 | 1.009220528 | 0.27096 | 3.724676461 | 0.000196 | 0.00642868 |
| ENSG00000261177.1 | 24.78307435 | 0.767677592 | 0.20725 | 3.704122645 | 0.000212 | 0.00684182 |
| ENSG00000279118.1 | 466.5558421 | 0.647297381 | 0.17517 | 3.695316519 | 0.00022 | 0.00699566 |
| ENSG00000136205.16 | 2962.642198 | 0.601070702 | 0.16282 | 3.691556283 | 0.000223 | 0.00707435 |
| ENSG00000143819.12 | 1609.617964 | 0.608374645 | 0.16503 | 3.686355542 | 0.000227 | 0.00709475 |
| ENSG00000162407.8 | 2750.830825 | 0.805592095 | 0.21861 | 3.685015315 | 0.000229 | 0.00709475 |
| ENSG00000183579.15 | 744.6443728 | 0.613809728 | 0.16646 | 3.687320536 | 0.000227 | 0.00709475 |
| ENSG00000203396.3 | 170.1716001 | 1.314275206 | 0.35668 | 3.684747499 | 0.000229 | 0.00709475 |
| ENSG00000100968.13 | 335.8640638 | 0.633103094 | 0.17188 | 3.683477188 | 0.00023 | 0.0071088 |
| ENSG00000204099.11 | 183.0439696 | 0.594732386 | 0.16162 | 3.679921202 | 0.000233 | 0.00716563 |
| ENSG00000127418.14 | 601.952864 | 0.708273108 | 0.19316 | 3.666860809 | 0.000246 | 0.00743419 |
| ENSG00000171903.16 | 129.3399371 | 0.745310493 | 0.20324 | 3.667102387 | 0.000245 | 0.00743419 |
| ENSG00000197496.5 | 114.8715109 | 0.702967648 | 0.19171 | 3.66673953 | 0.000246 | 0.00743419 |
| ENSG00000185338.4 | 9.129556243 | 0.988950786 | 0.27025 | 3.659350683 | 0.000253 | 0.00760639 |
| ENSG00000101850.12 | 77.92987865 | 0.649758547 | 0.17765 | 3.657479363 | 0.000255 | 0.00761828 |
| ENSG00000165584.15 | 9.342034198 | 0.749016226 | 0.20528 | 3.648817096 | 0.000263 | 0.00774828 |
| ENSG00000104760.16 | 17.3212411 | 0.862722525 | 0.23652 | 3.647513207 | 0.000265 | 0.00775669 |
| ENSG00000237887.1 | 40.0703658 | 0.632151217 | 0.17334 | 3.646955116 | 0.000265 | 0.00775669 |
| ENSG00000092820.17 | 2683.500805 | 0.646632447 | 0.1775 | 3.642901217 | 0.00027 | 0.00779138 |
| ENSG00000166183.15 | 22.95042329 | 0.630665087 | 0.17335 | 3.638088044 | 0.000275 | 0.00791618 |
| ENSG00000117834.12 | 18.65487822 | 0.680105689 | 0.18704 | 3.636093824 | 0.000277 | 0.0079554 |

| | | | | | | |
|---|---|---|---|---|---|---|
| ENSG00000074047.21 | 55.00242015 | 0.910099872 | 0.25044 | 3.633988922 | 0.000279 | 0.00799827 |
| ENSG00000254584.1 | 151.9211188 | 1.122493986 | 0.30922 | 3.630051172 | 0.000283 | 0.00809873 |
| ENSG00000141756.18 | 378.7999679 | 0.641283891 | 0.1767 | 3.629216033 | 0.000284 | 0.00810247 |
| ENSG00000016402.13 | 72.52589793 | 0.64615279 | 0.17846 | 3.620784315 | 0.000294 | 0.00830219 |
| ENSG00000133048.12 | 372.853493 | 0.864076082 | 0.23959 | 3.606409183 | 0.00031 | 0.00854494 |
| ENSG00000134873.9 | 550.9560192 | 0.763896807 | 0.21213 | 3.600997865 | 0.000317 | 0.00854494 |
| ENSG00000230062.5 | 15.63645612 | -0.707932879 | 0.19652 | -3.602358411 | 0.000315 | 0.00854494 |
| ENSG00000238099.2 | 12.21722061 | 0.858871956 | 0.23845 | 3.601963393 | 0.000316 | 0.00854494 |
| ENSG00000267534.2 | 294.2147278 | 1.101074242 | 0.30551 | 3.604107123 | 0.000313 | 0.00854494 |
| ENSG00000106571.12 | 241.3095255 | 0.833036944 | 0.23156 | 3.59747598 | 0.000321 | 0.00858481 |
| ENSG00000156076.9 | 1289.598346 | 0.87841316 | 0.2443 | 3.595597526 | 0.000324 | 0.00858481 |
| ENSG00000166819.11 | 77.0222583 | 0.659851856 | 0.18366 | 3.592857037 | 0.000327 | 0.00858481 |
| ENSG00000235989.3 | 39.5876589 | 0.732352774 | 0.20371 | 3.595123119 | 0.000324 | 0.00858481 |
| ENSG00000268038.1 | 36.32630445 | 0.917431736 | 0.25559 | 3.589424473 | 0.000331 | 0.00861077 |
| ENSG00000025423.11 | 241.2882263 | 0.655639082 | 0.18342 | 3.574453591 | 0.000351 | 0.00900506 |
| ENSG00000259479.6 | 42.88380684 | -1.413301746 | 0.39576 | -3.571112869 | 0.000355 | 0.00904193 |
| ENSG00000110436.11 | 59030.34413 | 0.799737892 | 0.2242 | 3.567019594 | 0.000361 | 0.00905216 |
| ENSG00000198417.6 | 280.0516054 | 0.604014981 | 0.16924 | 3.569087978 | 0.000358 | 0.00905216 |
| ENSG00000135744.7 | 2616.247664 | 0.716406042 | 0.20126 | 3.559581911 | 0.000371 | 0.00920917 |
| ENSG00000170412.16 | 165.4243534 | 0.653971789 | 0.18391 | 3.555993181 | 0.000377 | 0.00928541 |
| ENSG00000109113.18 | 318.8598528 | 0.6513732 | 0.18333 | 3.553062596 | 0.000381 | 0.00930249 |
| ENSG00000140459.17 | 36.721183 | 0.70594151 | 0.1987 | 3.552888982 | 0.000381 | 0.00930249 |
| ENSG00000261402.1 | 36.0021684 | -0.725223919 | 0.20414 | -3.552534666 | 0.000382 | 0.00930249 |
| ENSG00000134042.12 | 1277.187301 | 0.60457224 | 0.17097 | 3.536214951 | 0.000406 | 0.00964515 |
| ENSG00000278962.1 | 83.48360588 | -0.651837156 | 0.18437 | -3.535408015 | 0.000407 | 0.00965236 |
| ENSG00000125398.5 | 1523.246289 | 0.758175862 | 0.21487 | 3.528510792 | 0.000418 | 0.00982286 |
| ENSG00000125820.5 | 142.8644866 | 0.597050072 | 0.16946 | 3.523284901 | 0.000426 | 0.00998977 |
| ENSG00000141485.16 | 339.4684244 | 0.717993935 | 0.20424 | 3.515418374 | 0.000439 | 0.01012902 |
| ENSG00000152661.7 | 6850.648862 | 0.850352892 | 0.2419 | 3.515252167 | 0.000439 | 0.01012902 |
| ENSG00000180340.6 | 22.23337455 | 0.761934971 | 0.21678 | 3.514821392 | 0.00044 | 0.01012902 |
| ENSG00000225174.1 | 35.50946368 | 0.762370025 | 0.21671 | 3.517982434 | 0.000435 | 0.01012902 |
| ENSG00000161955.16 | 430.8286807 | 0.631261836 | 0.18029 | 3.501374122 | 0.000463 | 0.01042106 |
| ENSG00000079215.13 | 13631.38119 | 0.823374261 | 0.23522 | 3.500426148 | 0.000465 | 0.01043536 |
| ENSG00000133433.10 | 297.6652255 | -0.679755824 | 0.19445 | -3.495804692 | 0.000473 | 0.01046536 |
| ENSG00000164976.8 | 2055.755167 | 0.614518558 | 0.17573 | 3.496951388 | 0.000471 | 0.01046536 |
| ENSG00000241839.9 | 665.4163313 | 0.708607736 | 0.20267 | 3.496391859 | 0.000472 | 0.01046536 |
| ENSG00000114790.12 | 791.5308848 | 0.588255609 | 0.16857 | 3.48961298 | 0.000484 | 0.01058936 |
| ENSG00000169860.6 | 299.9167946 | 0.708550246 | 0.20299 | 3.490560054 | 0.000482 | 0.01058936 |
| ENSG00000184232.8 | 657.7467697 | 0.661321787 | 0.1895 | 3.489792446 | 0.000483 | 0.01058936 |

| | | | | | | |
|---|---|---|---|---|---|---|
| ENSG00000124145.6 | 932.3735459 | 0.940391012 | 0.27047 | 3.476885539 | 0.000507 | 0.01096501 |
| ENSG00000135540.11 | 465.2531474 | 0.636690313 | 0.1834 | 3.4715573 | 0.000517 | 0.01109173 |
| ENSG00000178602.7 | 11.83954008 | 0.935318439 | 0.26984 | 3.466226816 | 0.000528 | 0.01117432 |
| ENSG00000166148.3 | 43.29560881 | -0.748151118 | 0.21596 | -3.464245016 | 0.000532 | 0.01123385 |
| ENSG00000129151.8 | 241.9193087 | 0.695004429 | 0.20067 | 3.463468421 | 0.000533 | 0.01124325 |
| ENSG00000248394.1 | 12.01656905 | 0.597105765 | 0.17261 | 3.459199181 | 0.000542 | 0.01135319 |
| ENSG00000219607.3 | 150.6593163 | 0.638188752 | 0.18521 | 3.445790951 | 0.000569 | 0.01162324 |
| ENSG00000250602.5 | 34.01684249 | 0.629337301 | 0.1828 | 3.442770279 | 0.000576 | 0.01168417 |
| ENSG00000214954.8 | 213.9119918 | 1.128905008 | 0.32857 | 3.435761283 | 0.000591 | 0.01187438 |
| ENSG00000225630.1 | 2626.084187 | 1.916272249 | 0.55841 | 3.431656122 | 0.0006 | 0.01195026 |
| ENSG00000054179.11 | 275.4626886 | 0.671268286 | 0.19625 | 3.420545011 | 0.000625 | 0.01231838 |
| ENSG00000155324.9 | 1094.31369 | 0.596748973 | 0.17452 | 3.419340074 | 0.000628 | 0.01232583 |
| ENSG00000144230.16 | 404.3460183 | 0.648402422 | 0.18986 | 3.41521916 | 0.000637 | 0.01249001 |
| ENSG00000184608.8 | 10.51317746 | 0.792761574 | 0.23225 | 3.413354535 | 0.000642 | 0.01250203 |
| ENSG00000180613.10 | 8.932333956 | 0.824539855 | 0.2433 | 3.388953558 | 0.000702 | 0.01324542 |
| ENSG00000257434.1 | 683.9654209 | -0.613439962 | 0.18112 | -3.386925253 | 0.000707 | 0.01331929 |
| ENSG00000169856.8 | 10.27342817 | 0.641010734 | 0.1895 | 3.382579492 | 0.000718 | 0.01340903 |
| ENSG00000106565.17 | 210.651003 | 0.654778108 | 0.19405 | 3.37433982 | 0.00074 | 0.01374212 |
| ENSG00000162621.6 | 78.76876661 | -0.616398993 | 0.18322 | -3.36428096 | 0.000767 | 0.01410022 |
| ENSG00000236953.1 | 59.97226322 | 1.096655002 | 0.32596 | 3.364396342 | 0.000767 | 0.01410022 |
| ENSG00000259984.1 | 11.80162955 | -0.61262572 | 0.18236 | -3.359346673 | 0.000781 | 0.01417722 |
| ENSG00000179604.9 | 1578.771421 | 0.612035961 | 0.18256 | 3.352566582 | 0.000801 | 0.01440208 |
| ENSG00000164188.8 | 802.7233254 | 0.729068456 | 0.21775 | 3.348131494 | 0.000814 | 0.0145582 |
| ENSG00000172201.11 | 1299.697685 | 0.614682141 | 0.1837 | 3.346063538 | 0.00082 | 0.01458434 |
| ENSG00000267714.1 | 15.58186604 | 0.728583228 | 0.2178 | 3.345172455 | 0.000822 | 0.01458434 |
| ENSG00000134250.18 | 1322.903957 | 0.681038483 | 0.20402 | 3.338114684 | 0.000843 | 0.01473457 |
| ENSG00000181284.2 | 17.71968742 | 0.587120946 | 0.17595 | 3.336789394 | 0.000848 | 0.01473807 |
| ENSG00000225472.1 | 54.64097701 | 0.586551554 | 0.17607 | 3.331286195 | 0.000864 | 0.01494856 |
| ENSG00000242252.1 | 19.68025147 | 0.620911435 | 0.18638 | 3.331443476 | 0.000864 | 0.01494856 |
| ENSG00000131721.5 | 12.43737413 | 0.684837201 | 0.20619 | 3.321450184 | 0.000896 | 0.01521245 |
| ENSG00000270181.1 | 150.4931574 | -0.69450189 | 0.20989 | -3.308946888 | 0.000936 | 0.01561136 |
| ENSG00000136869.13 | 422.238597 | 0.595860179 | 0.1803 | 3.304878075 | 0.00095 | 0.01573628 |
| ENSG00000270118.1 | 28.24745898 | 0.848326334 | 0.25675 | 3.30409568 | 0.000953 | 0.01573628 |
| ENSG00000232987.1 | 10.51580287 | 0.663922409 | 0.20109 | 3.301580669 | 0.000961 | 0.01585259 |
| ENSG00000275395.4 | 152.9578004 | -0.805713367 | 0.2448 | -3.291276669 | 0.000997 | 0.0161346 |
| ENSG00000163576.17 | 58.88986593 | 0.643330989 | 0.19553 | 3.290245037 | 0.001001 | 0.01616846 |
| ENSG00000148482.11 | 659.0559597 | 0.819634372 | 0.24947 | 3.285523482 | 0.001018 | 0.01639043 |
| ENSG00000143772.9 | 2529.253101 | 0.638430515 | 0.19443 | 3.283579202 | 0.001025 | 0.0164268 |
| ENSG00000259895.1 | 520.2574972 | 1.123641519 | 0.34346 | 3.271503613 | 0.00107 | 0.01680441 |

| | | | | | | |
|---|---|---|---|---|---|---|
| ENSG00000218109.5 | 11.29592545 | -0.76343552 | 0.23418 | -3.260079982 | 0.001114 | 0.01715564 |
| ENSG00000215612.7 | 30.72959004 | 0.607930977 | 0.18666 | 3.256871223 | 0.001126 | 0.01727318 |
| ENSG00000151012.13 | 1785.418461 | 0.618186401 | 0.19006 | 3.252573191 | 0.001144 | 0.01748444 |
| ENSG00000167880.7 | 33.09897763 | -0.660487541 | 0.20333 | -3.248381743 | 0.001161 | 0.01766534 |
| ENSG00000254910.1 | 12.11718546 | 0.831050465 | 0.25583 | 3.248503099 | 0.00116 | 0.01766534 |
| ENSG00000168306.12 | 69.98072529 | 0.588580048 | 0.18155 | 3.241927672 | 0.001187 | 0.0178821 |
| ENSG00000275620.1 | 756.1592185 | 0.614970829 | 0.18973 | 3.241298185 | 0.00119 | 0.0178821 |
| ENSG00000080493.14 | 3044.361722 | 0.73794046 | 0.2278 | 3.239431117 | 0.001198 | 0.01793737 |
| ENSG00000229847.8 | 1687.91865 | 0.603900216 | 0.18672 | 3.234234195 | 0.00122 | 0.01813502 |
| ENSG00000236523.2 | 203.9502731 | 1.198672785 | 0.37055 | 3.234804305 | 0.001217 | 0.01813502 |
| ENSG00000164708.5 | 47.22939984 | 0.6488903 | 0.20069 | 3.233248921 | 0.001224 | 0.01817142 |
| ENSG00000275763.3 | 12.63058699 | 0.662941984 | 0.20544 | 3.226987454 | 0.001251 | 0.01841436 |
| ENSG00000216285.5 | 11.46404363 | -0.87028065 | 0.27004 | -3.222824737 | 0.001269 | 0.01865737 |
| ENSG00000121764.11 | 18.70900352 | -0.675953612 | 0.20999 | -3.218939736 | 0.001287 | 0.01880454 |
| ENSG00000072952.18 | 1491.412366 | 0.657591892 | 0.20533 | 3.202614615 | 0.001362 | 0.0194948 |
| ENSG00000255337.1 | 12.16490408 | 0.627212759 | 0.19579 | 3.203418836 | 0.001358 | 0.0194948 |
| ENSG00000261195.1 | 16.47587705 | 0.654278923 | 0.20469 | 3.196418899 | 0.001391 | 0.01969274 |
| ENSG00000164199.15 | 2954.916236 | 0.60883477 | 0.19126 | 3.183277066 | 0.001456 | 0.02006127 |
| ENSG00000186081.11 | 70.09056204 | -0.64841967 | 0.20391 | -3.179890196 | 0.001473 | 0.02013018 |
| ENSG00000233273.1 | 8.861566175 | -0.712405891 | 0.22459 | -3.172089785 | 0.001513 | 0.02034593 |
| ENSG00000189334.8 | 9.062268824 | 0.749185889 | 0.23743 | 3.155415611 | 0.001603 | 0.02101366 |
| ENSG00000276085.1 | 56.08570744 | -0.782801701 | 0.24816 | -3.154453131 | 0.001608 | 0.02103731 |
| ENSG00000104921.14 | 9.75861104 | -0.93532449 | 0.29782 | -3.140621942 | 0.001686 | 0.02168272 |
| ENSG00000136160.14 | 1638.146173 | 0.758519289 | 0.2417 | 3.138243533 | 0.0017 | 0.02172367 |
| ENSG00000251165.5 | 25.44422622 | 0.692751037 | 0.22069 | 3.139085787 | 0.001695 | 0.02172367 |
| ENSG00000007129.17 | 8.958606583 | -0.747872766 | 0.23872 | -3.13289575 | 0.001731 | 0.02187558 |
| ENSG00000278996.1 | 33.18241642 | 0.636764113 | 0.20365 | 3.126735004 | 0.001768 | 0.02215195 |
| ENSG00000280441.2 | 33.18241642 | 0.636764113 | 0.20365 | 3.126735004 | 0.001768 | 0.02215195 |
| ENSG00000279778.1 | 20.88641557 | -0.627887216 | 0.20136 | -3.118300048 | 0.001819 | 0.02253524 |
| ENSG00000233139.1 | 231.6763465 | 1.1757812 | 0.37777 | 3.112412866 | 0.001856 | 0.02262175 |
| ENSG00000270001.1 | 22.71473889 | 0.817118568 | 0.26306 | 3.106150642 | 0.001895 | 0.02291631 |
| ENSG00000188818.12 | 367.8559931 | -0.600598619 | 0.19349 | -3.104075071 | 0.001909 | 0.02299097 |
| ENSG00000228412.7 | 28.46849809 | 0.60670058 | 0.19556 | 3.102331873 | 0.00192 | 0.02302464 |
| ENSG00000259605.3 | 14.98590301 | 0.707654083 | 0.22822 | 3.100785283 | 0.00193 | 0.02311829 |
| ENSG00000225938.1 | 14.61604224 | 0.793410671 | 0.25612 | 3.097815849 | 0.00195 | 0.02316243 |
| ENSG00000233435.2 | 17.35060523 | -0.705222161 | 0.22864 | -3.084417033 | 0.00204 | 0.02365791 |
| ENSG00000232035.1 | 1842.695111 | 1.235235814 | 0.40057 | 3.083658361 | 0.002045 | 0.02369161 |
| ENSG00000242221.8 | 845.9835976 | 1.51959177 | 0.49319 | 3.081178786 | 0.002062 | 0.02386293 |
| ENSG00000187193.8 | 441.989164 | 0.930677626 | 0.3032 | 3.069566669 | 0.002144 | 0.0243449 |

| | | | | | | |
|---|---|---|---|---|---|---|
| ENSG00000119938.8 | 981.2739234 | 0.704683057 | 0.23111 | 3.049144779 | 0.002295 | 0.02540884 |
| ENSG00000184414.2 | 123.1835998 | 1.283680394 | 0.42167 | 3.044283078 | 0.002332 | 0.02566589 |
| ENSG00000157654.17 | 525.5944683 | -0.627594139 | 0.20792 | -3.018500387 | 0.00254 | 0.02686233 |
| ENSG00000210194.1 | 15.66046073 | 0.926131114 | 0.30725 | 3.014257237 | 0.002576 | 0.0269931 |
| ENSG00000223774.5 | 20.77568056 | 0.612563872 | 0.20335 | 3.012393352 | 0.002592 | 0.02705632 |
| ENSG00000188662.6 | 11.98831742 | 0.632337685 | 0.21017 | 3.008707155 | 0.002624 | 0.02721211 |
| ENSG00000259446.5 | 10.01645383 | 0.91348855 | 0.30405 | 3.004363824 | 0.002661 | 0.0274301 |
| ENSG00000007908.15 | 23.53904631 | -1.147124276 | 0.38197 | -3.003185549 | 0.002672 | 0.02746161 |
| ENSG00000143851.15 | 35.88599994 | -0.667813933 | 0.22366 | -2.985798125 | 0.002828 | 0.02803597 |
| ENSG00000255390.1 | 28.87218927 | 0.871677909 | 0.29338 | 2.971145537 | 0.002967 | 0.02883934 |
| ENSG00000241231.1 | 71.57019891 | 0.721509069 | 0.24338 | 2.964526034 | 0.003031 | 0.02905924 |
| ENSG00000111432.4 | 12.83850631 | 0.733537521 | 0.24795 | 2.958363183 | 0.003093 | 0.02930162 |
| ENSG00000264617.1 | 8.871917158 | -0.785687904 | 0.26569 | -2.957134965 | 0.003105 | 0.02933755 |
| ENSG00000183242.11 | 134.5916676 | 1.223879284 | 0.41413 | 2.955334076 | 0.003123 | 0.02940137 |
| ENSG00000225972.1 | 2785.403504 | -1.835374633 | 0.62194 | -2.951034354 | 0.003167 | 0.02967802 |
| ENSG00000279082.3 | 38.51003777 | 0.625326514 | 0.21252 | 2.942498777 | 0.003256 | 0.03031531 |
| ENSG00000147145.12 | 738.5034729 | 1.284273089 | 0.43795 | 2.932449907 | 0.003363 | 0.03100522 |
| ENSG00000270136.5 | 66.1684677 | 0.75121979 | 0.25986 | 2.890812565 | 0.003842 | 0.0332508 |
| ENSG00000197901.11 | 43.02357169 | 0.639148189 | 0.2214 | 2.886875204 | 0.003891 | 0.03346909 |
| ENSG00000188133.5 | 52.68972372 | -0.609776207 | 0.21232 | -2.87199923 | 0.004079 | 0.03428456 |
| ENSG00000115602.16 | 52.38157609 | 1.289688681 | 0.44978 | 2.867389445 | 0.004139 | 0.03448998 |
| ENSG00000261325.1 | 44.05721438 | -0.621288135 | 0.21797 | -2.850272514 | 0.004368 | 0.035529 |
| ENSG00000197632.8 | 10.36795536 | -0.613650577 | 0.21538 | -2.849184532 | 0.004383 | 0.03555336 |
| ENSG00000157335.20 | 17.00825929 | 0.743135647 | 0.2623 | 2.833196899 | 0.004608 | 0.03650258 |
| ENSG00000132470.13 | 537.8448647 | 0.649716331 | 0.22937 | 2.832571034 | 0.004618 | 0.03651789 |
| ENSG00000162877.12 | 23.53562487 | 0.742527888 | 0.26222 | 2.831671207 | 0.004631 | 0.03659268 |
| ENSG00000279123.1 | 8.960787213 | 0.679488495 | 0.2408 | 2.821777158 | 0.004776 | 0.03736024 |
| ENSG00000229206.3 | 49.34496978 | 0.665791119 | 0.23706 | 2.808542567 | 0.004977 | 0.03815047 |
| ENSG00000104415.13 | 65.16243061 | -0.678099405 | 0.24186 | -2.80367143 | 0.005052 | 0.03830841 |
| ENSG00000267056.2 | 19.05726843 | 0.783891222 | 0.2815 | 2.784670243 | 0.005358 | 0.03963387 |
| ENSG00000139155.8 | 742.510556 | 0.606623673 | 0.21911 | 2.768627209 | 0.005629 | 0.04063536 |
| ENSG00000265015.1 | 9.952431653 | -0.586978492 | 0.2121 | -2.767470861 | 0.005649 | 0.04067582 |
| ENSG00000211896.7 | 19.10356377 | -1.322391834 | 0.47836 | -2.764439263 | 0.005702 | 0.04094112 |
| ENSG00000155307.17 | 50.68880063 | -0.759689906 | 0.27553 | -2.757181852 | 0.00583 | 0.04134172 |
| ENSG00000182508.13 | 42.31100657 | 0.663092406 | 0.24072 | 2.754611772 | 0.005876 | 0.04149625 |
| ENSG00000240590.1 | 454.6793493 | 1.326889102 | 0.48367 | 2.743354505 | 0.006081 | 0.04222169 |
| ENSG00000205364.3 | 137.2433497 | 0.688223906 | 0.25095 | 2.742445084 | 0.006098 | 0.04231023 |
| ENSG00000277734.6 | 34.0583155 | -0.641194994 | 0.23402 | -2.739909502 | 0.006146 | 0.04246621 |
| ENSG00000271127.1 | 27.41451297 | -0.630069954 | 0.23003 | -2.739113828 | 0.006161 | 0.04253642 |

| | | | | | | |
|---|---|---|---|---|---|---|
| ENSG00000132967.9 | 62.47656948 | -1.114175715 | 0.40691 | -2.738106999 | 0.006179 | 0.04258527 |
| ENSG00000198125.12 | 9.333600996 | 0.62255402 | 0.22959 | 2.711618209 | 0.006696 | 0.04458944 |
| ENSG00000118785.13 | 1139.542643 | -0.668326043 | 0.24713 | -2.704310352 | 0.006845 | 0.04497653 |
| ENSG00000141469.16 | 349.3141952 | 0.820644021 | 0.30537 | 2.687415023 | 0.007201 | 0.04642133 |
| ENSG00000185710.9 | 76.51759738 | -0.672377853 | 0.25143 | -2.674244189 | 0.00749 | 0.04739388 |
| ENSG00000266573.5 | 11.86523166 | -0.593136623 | 0.22223 | -2.669016719 | 0.007607 | 0.04781844 |
| ENSG00000258647.5 | 9.949154698 | -0.788107816 | 0.29596 | -2.662902357 | 0.007747 | 0.04835932 |
| ENSG00000248751.6 | 19.27273076 | 0.681644085 | 0.25745 | 2.64772163 | 0.008104 | 0.04965706 |

Abbreviations: baseMean, normalized read counts of all samples; log2Foldchange, effect size estimate; lfcSE, standard error of the logfoldchange; stat, Wald statistical test values; pvalue, uncorrected p-value; p-adj, corrected p-value

**Supplementary Table 3**

**Controls > Bipolar Disorder Simple Comparison**

| Ensemble ID | baseMean | log2FoldChange | lfcSE | stat | pvalue | padj |
|---|---|---|---|---|---|---|
| ENSG00000124440.15 | 618.85673 | 0.963640201 | 0.1918236 | 5.0235759 | 5.07E-07 | 0.000884 |
| ENSG00000187193.8 | 681.51935 | 1.761186273 | 0.3465509 | 5.0820427 | 3.73E-07 | 0.000884 |
| ENSG00000225972.1 | 2253.0391 | -2.832747638 | 0.5482977 | -5.166441 | 2.39E-07 | 0.000884 |
| ENSG00000155380.11 | 741.70462 | 0.618686889 | 0.140914 | 4.3905296 | 1.13E-05 | 0.007387 |
| ENSG00000167772.11 | 436.92392 | 1.728915437 | 0.3902062 | 4.4307743 | 9.39E-06 | 0.007387 |
| ENSG00000198886.2 | 482968.54 | 0.707747893 | 0.1577614 | 4.4861918 | 7.25E-06 | 0.007387 |
| ENSG00000239282.7 | 134.26168 | 0.668837237 | 0.1535617 | 4.3554944 | 1.33E-05 | 0.007709 |
| ENSG00000168394.10 | 316.70671 | 0.598704662 | 0.1413579 | 4.2353829 | 2.28E-05 | 0.009936 |
| ENSG00000133048.12 | 428.51157 | 1.141468155 | 0.2782615 | 4.1021412 | 4.09E-05 | 0.014763 |
| ENSG00000138772.12 | 160.14953 | 0.655455502 | 0.1606837 | 4.0791654 | 4.52E-05 | 0.014763 |
| ENSG00000164292.12 | 2138.929 | 0.612705959 | 0.1558793 | 3.9306445 | 8.47E-05 | 0.023316 |
| ENSG00000168209.4 | 726.33326 | 0.722729665 | 0.187128 | 3.8622206 | 0.000112 | 0.024184 |
| ENSG00000214456.8 | 173.77357 | 0.621219435 | 0.1610594 | 3.8570821 | 0.000115 | 0.024184 |
| ENSG00000125144.13 | 515.02886 | 0.839840643 | 0.2210446 | 3.7994165 | 0.000145 | 0.024821 |
| ENSG00000163395.16 | 154.64362 | 0.765116394 | 0.2019387 | 3.7888545 | 0.000151 | 0.024821 |
| ENSG00000196136.16 | 139.16433 | 2.19111007 | 0.58956 | 3.7165177 | 0.000202 | 0.030037 |
| ENSG00000205364.3 | 165.18001 | 1.084882983 | 0.2929361 | 3.7034797 | 0.000213 | 0.030037 |
| ENSG00000254245.2 | 217.19099 | 0.716294576 | 0.1960874 | 3.6529344 | 0.000259 | 0.033046 |
| ENSG00000136235.15 | 571.74489 | 0.743646179 | 0.2051343 | 3.625167 | 0.000289 | 0.035932 |
| ENSG00000225630.1 | 518.85726 | 0.916401344 | 0.2545259 | 3.6004249 | 0.000318 | 0.037734 |
| ENSG00000078401.6 | 207.03702 | 0.809790005 | 0.2271126 | 3.5655884 | 0.000363 | 0.038871 |
| ENSG00000212907.2 | 39665.086 | 0.608061415 | 0.1729062 | 3.516712 | 0.000437 | 0.040977 |
| ENSG00000185201.16 | 241.0811 | 0.840441354 | 0.2428545 | 3.4606786 | 0.000539 | 0.048549 |

Abbreviations: baseMean, normalized read counts of all samples; log2Foldchange, effect size estimate; lfcSE, standard error of the logfoldchange; stat, Wald statistical test values; pvalue, uncorrected p-value; p-adj, corrected p-value.
